# Transcription Factor Binding Site Prediction in Microbial Biosynthetic Gene Cluster Families

**DOI:** 10.64898/2026.11.24.768815

**Authors:** Idris Matine, Fernando Garcia Guevara, Frederick Clasen, Miao Guo, Saeed Shoaie

## Abstract

Transcriptional regulation of biosynthetic gene clusters (BGCs) determines whether secondary metabolites are produced, yet the regulatory elements controlling BGC expression remain largely uncharacterised. We present a comparative-genomic pipeline for systematic *de novo* discovery of conserved transcription factor (TF) binding sites across BGC families catalogued in the BiG-FAM database. Within each family, BGC-encoded genes were partitioned into clusters of orthologous genes, their upstream regions extracted and redundancy-filtered, and motif discovery used to identify sequences conserved within each cluster. Applied to 994 families (over 660,000 BGCs), the pipeline recovered experimentally characterised operators, including the EctR1 operator of ectoine clusters and both the Fur and IdeR iron boxes within a single siderophore family, each confined to the lineage in which its regulator operates. Annotation of BGC-encoded proteins against curated TF family profiles mapped pathway-specific regulators across 921 BGC families, identifying LysR as a generalist present in every biosynthetic class and class-restricted families such as LytTR in RiPP clusters. Sequence similarity networks of 20 regulator families, combined with clustering of their upstream motifs, resolved TF subfamilies whose conserved motifs match characterised binding sites. Together, the pipeline and its searchable resource provide a systematic means of prioritising candidate TF binding site relationships for experimental testing.

## INTRODUCTION

Microbial secondary metabolites (SMs), including antibiotics, immunosuppressants and anticancer agents, underpin major advances in medicine, agriculture and biotechnology (1–3). Their biosynthesis is typically encoded by biosynthetic gene clusters (BGCs), co-localised genomic regions containing the genes for metabolite assembly, regulation, transport and self-resistance (2, 4). Specialised computational tools have transformed the discovery and systematic analysis of BGCs antiSMASH enables genome-wide identification and annotation of BGCs (5), BiG-SCAPE and BiG-SLiCE group them into gene cluster families (GCFs) by sequence similarity (6), and the BiG-FAM database provides a pre-computed catalogue of GCFs spanning more than 1.2 million BGCs from bacterial, fungal and archaeal genomes (7).

The rapid expansion of microbial genomic data has revealed a largely untapped reservoir of biosynthetic potential. However, most BGCs remain transcriptionally silent under standard laboratory conditions, rendering the metabolites they encode inaccessible (8, 9). This widespread silencing is a regulatory barrier. Without understanding the transcriptional mechanisms that govern BGC expression, the biosynthetic potential revealed by genome mining cannot be translated into the discovery of new bioactive compounds.

Transcription factors (TFs) regulating BGCs fall into two broad classes (10). Pathway-specific transcription factors (PSTFs) are encoded within or adjacent to BGCs and control cluster expression directly by binding operator sequences in upstream promoter regions, as exemplified by the actinorhodin activator ActII-ORF4 of *Streptomyces coelicolor* (11, 12). Global transcription factors (GTFs) regulate broader cellular physiology and influence secondary metabolism indirectly (13, 14). Mapping transcription factor binding sites (TFBSs) in BGC promoter regions is therefore central to understanding whether a pathway remains silent or becomes active.

Experimental mapping of TFBSs is laborious and inherently hypothesis-driven, making systematic characterisation across the rapidly expanding BGC repertoire impractical, and computational approaches have become essential for scaling regulatory inference. Two broad strategies exist. Motif-scanning approaches search promoter regions with position weight matrices (PWMs) of characterised TFs. The TFBS Finder module of antiSMASH applies this to individual clusters, annotating candidate sites within cluster-associated promoters (15). COMMBAT (COnditions for Microbial Metabolite Biosynthesis Activated Transcription) extends the approach by integrating PWM matching with genomic context and gene-function information in a unified scoring framework, improving sensitivity to the short, degenerate and potentially low-affinity sites characteristic of many BGC regulatory regions (16). These approaches prioritise regulatory hypotheses, culture conditions and engineering targets, but they are restricted to TFs whose binding preferences have already been characterised and cannot identify the binding sites of the many BGC-associated regulators for which no PWM exists. *De novo* motif discovery tools such as the MEME Suite avoid this dependence by identifying over-represented sequence patterns within a set of putatively co-regulated upstream regions without prior knowledge of the cognate TF (17).

The AURTHO framework (AUtoregulation of oRTHOlogous transcription factors) defines such a set from orthology. Because TFs frequently autoregulate, comparing the upstream regions of orthologous LacI TFs across species yielded substantially more TFBS predictions (18). The same comparative logic is not restricted to autoregulation. Orthologous genes performing the same function within a conserved pathway are likely to share regulatory control, so their upstream regions form a natural input for de novo discovery irrespective of which TF binds them. GCFs provide exactly this structure, since every biosynthetic, transport and regulatory gene in a family has orthologues across many producer organisms. To our knowledge, this orthology-guided strategy has not been applied systematically to BGCs.

Here, we present a pipeline that combines BiG-FAM GCF assignments with orthologous gene grouping, upstream sequence extraction, redundancy filtering and de novo motif discovery, and links the resulting motifs to candidate regulators through HMM-based annotation of BGC-encoded TFs. We apply it to 994 GCFs, describe the distribution of BGC-encoded TF families across biosynthetic classes, test whether motifs discovered without reference to known sites recover experimentally characterised operators, and relate conserved motifs to TF sequence subfamilies within major regulator families. The pipeline is openly available, and all predicted motifs and TF annotations are provided through a searchable public resource.

## METHODS

### Data

All BGCs analysed in this study were obtained from the BiG-FAM database(7), which contains nearly 1.2 million BGCs from bacterial, fungal and archaeal genomes grouped into 138,345 GCFs on the basis of protein-domain similarity using BiG-SLiCE(19). Because *de novo* motif discovery requires multiple diverse sequences per group, GCFs were filtered to retain only those with at least 100 member BGCs (1,263 families). GCF identifiers (FAM####) throughout this study are the BiG-FAM family IDs.

### Pipeline implementation

The complete workflow was implemented in Nextflow (v23.10.1) (20). All third-party bioinformatics tools were executed within Singularity containers pulled from public BioContainers and Docker repositories. The data-handling, parsing and analysis steps that connect these tools were implemented as custom Python v3.9 modules.

### Orthologous gene construction

For each GCF, the protein sequences of all genes encoded by its member BGCs were compared all-versus-all with PROTEINORTHO (v6.3.5) with the DIAMOND alignment backend and default parameters (21, 22). PROTEINORTHO partitions the reciprocal-best-hit graph by spectral clustering into groups of orthologous genes, here termed clusters of orthologous genes (COGs). COGs are defined within each GCF and are unrelated to the NCBI COG database.

### Upstream region extraction

For each gene assigned to a COG, the non-coding region upstream of the predicted translation start site was extracted from the BGC region sequence. Extraction was bounded to a maximum of 350 bp and a minimum of 100 bp upstream of the start codon, with an additional 20 bp of coding sequence included to capture sites overlapping the translation start. These boundaries were selected on the basis of a prior meta-analysis of BGC promoter architecture and experimentally characterised TFBS positions within biosynthetic clusters, which indicated that most functional binding sites fall within this window (10).

#### Redundancy reduction

Upstream regions from closely related strains or near-identical BGC copies would otherwise dominate motif discovery. Within each COG, extracted regions were clustered with CD-HIT-EST (v4.8.1)(23), at 90% nucleotide identity (-c 0.90 -n 5 -d 0) and only the representative sequence of each cluster was retained.

#### Motif discovery

Motif discovery was performed with MEME (MEME Suite v5.5.8) (17) on the dereplicated upstream sequences of each COG with at least four such sequences. Six runs were executed per COG, combining two occurrence models, ZOOPS (zero or one occurrence per sequence, corresponding to canonical single-site binding) and ANR (any number of repetitions), with maximum motif widths of 10, 20 and 30 bp to span the size range of bacterial TFBSs. All runs searched both strands (-revcomp) and reported up to 10 motifs. Motifs with E ≤ 0.05 were considered significant and were ranked by E-value within each COG. Unless otherwise stated, analyses use ZOOPS with a maximum width of 30 bp (zoops_maxw30).

### Annotation of BGC-encoded transcription factors

TF family profiles were built from ENTRAF (24), a curated collection of 1,784 experimentally characterised bacterial TFs annotated by Pfam domain family. Sequences of each family were aligned with MAFFT (v7.526, --auto) (25), and a profile HMM was built with HMMBUILD (HMMER v3.4) (26), yielding 190 family-specific profiles. The profiles were pressed into a single database with HMMPRESS and searched with HMMSEARCH (E ≤ 1×10⁻¹⁰, best hit per gene) against the 7.5 million non-redundant protein sequences encoded by all BiG-FAM BGCs.

Seven ENTRAF families whose defining domain is enzymatic rather than DNA-binding (GDXG, class-I PLP-dependent aminotransferase, protein kinase, adh_short_C2, cytochrome P450, PutA and acyl-CoA dehydrogenase) were excluded from TF annotation.

Hits were mapped to their COGs, and a COG was designated a TF COG when at least 90% of its members (minimum five) received a TF family annotation. TF COGs are treated as PSTFs, i.e. regulators encoded within the cluster, without implying that each regulates only its own cluster. For each GCF, the dominant PSTF was defined as the TF family of the TF COG present in the largest fraction of the family’s member BGCs.

### Recovery of experimentally characterised binding sites

To test whether *de novo* motifs recover known regulatory elements, GCFs containing MIBiG-curated BGCs with experimentally characterised operators were identified. For each family, a reference PWM was built from the published operator or consensus sequence and compared with TOMTOM (Pearson distance, default parameters) against all significant motifs predicted upstream of COGs in the family. Motifs from all six MEME settings were searched. Matches were considered significant at q ≤ 0.05. Genes co-transcribed with the operator-carrying gene served as within-cluster negative controls.

### Sequence similarity network analysis of TF families

To relate TF sequence subfamilies to their upstream motifs, sequence similarity networks (SSNs) were built for the 20 most prevalent PSTF families, each combining pipeline-annotated TF sequences with experimentally characterised regulators of the same family. Four are presented in detail: LysR and AraC/XylS (among the most prevalent), LytTR (the most class-restricted, to RiPPs) and MarR (the family of EctR1).

Characterised regulators and their binding-site matrices were obtained from PRODORIC (27), and their UniProt sequences were assigned to ENTRAF families with HMMSEARCH (E ≤ 1 × 10⁻¹⁰). EctR1 (UniProt G4T4A5), which is absent from PRODORIC, was added to the MarR network with the PWM used in the recovery analysis (28, 29).

For each family, pipeline-derived TF sequences were deduplicated at 90% identity with CD-HIT, combined with all family-assigned PRODORIC sequences, and compared all-by-all with EFI-EST (30). Each network was thresholded at an alignment score chosen from its percent-identity versus alignment-score plot (**Supplementary Table S1**) and clustered with MCL (inflation 2.0) (31), to define TF subfamilies (SSN TF clusters).

For each SSN TF cluster, motifs from all member GCFs (zoops_maxw30, MEME E ≤ 0.001, stricter than the catalogue threshold because motifs are pooled across families) were compared all-against-all with TOMTOM (Pearson distance, minimum overlap 5 bp). Pairs with q ≤ 0.1 formed a motif similarity network with edges weighted by −log₁₀(q), which was clustered with MCL (inflation 2.0). This edge threshold is permissive because single edges are not interpreted: clusters require groups of mutually similar motifs that recur across families. Motif clusters contributed by at least half of the GCFs in the parent SSN TF cluster (family coverage ≥ 0.5) were designated conserved. Where an SSN TF cluster contained a PRODORIC TF with a PWM, conserved motifs were compared against it with TOMTOM. Matches were considered significant at q ≤ 0.05 and matches with 0.05 < q ≤ 0.1 are reported as borderline. Networks were visualised in Cytoscape (32).

### AlphaFold3 structural modelling

TF–DNA complexes were modelled with the AlphaFold3 server (33).Each TF was modelled as a monomer and as a homodimer, alone and together with a double-stranded DNA fragment corresponding to a predicted motif (forward strand plus reverse complement), and in parallel with a composition-matched control generated by shuffling the motif sequence with a fixed random seed (seed = 42). The interface predicted TM-score (ipTM) was used as the confidence measure and pTM as a secondary indicator of fold quality.

### Statistical analyses

Motif significance is the MEME E-value, the expected number of motifs with equal or greater log-likelihood ratio in shuffled input sequences. TOMTOM q-values are Benjamini–Hochberg corrected. The relationship between COG conservation and the number of significant motifs was assessed with Spearman’s rank correlation across unbinned COGs. Upstream intergenic distances of TF and non-TF genes were compared with a two-sided Mann–Whitney U test at the gene level and at the level of per-COG medians. P-values below the floating-point limit are reported as P < 10⁻¹⁵.

## RESULTS

### Pipeline overview and dataset statistics

The pipeline takes the member BGCs of a GCF, partitions their encoded proteins into COGs, extracts the upstream region of each gene, and performs de novo motif discovery to identify conserved TFBSs shared by orthologous genes. In parallel, HMM profiles built from ENTRAF assign BGC-encoded genes to TF families, linking predicted sites to candidate regulators (**Figure 1A**).

**Figure 1:**
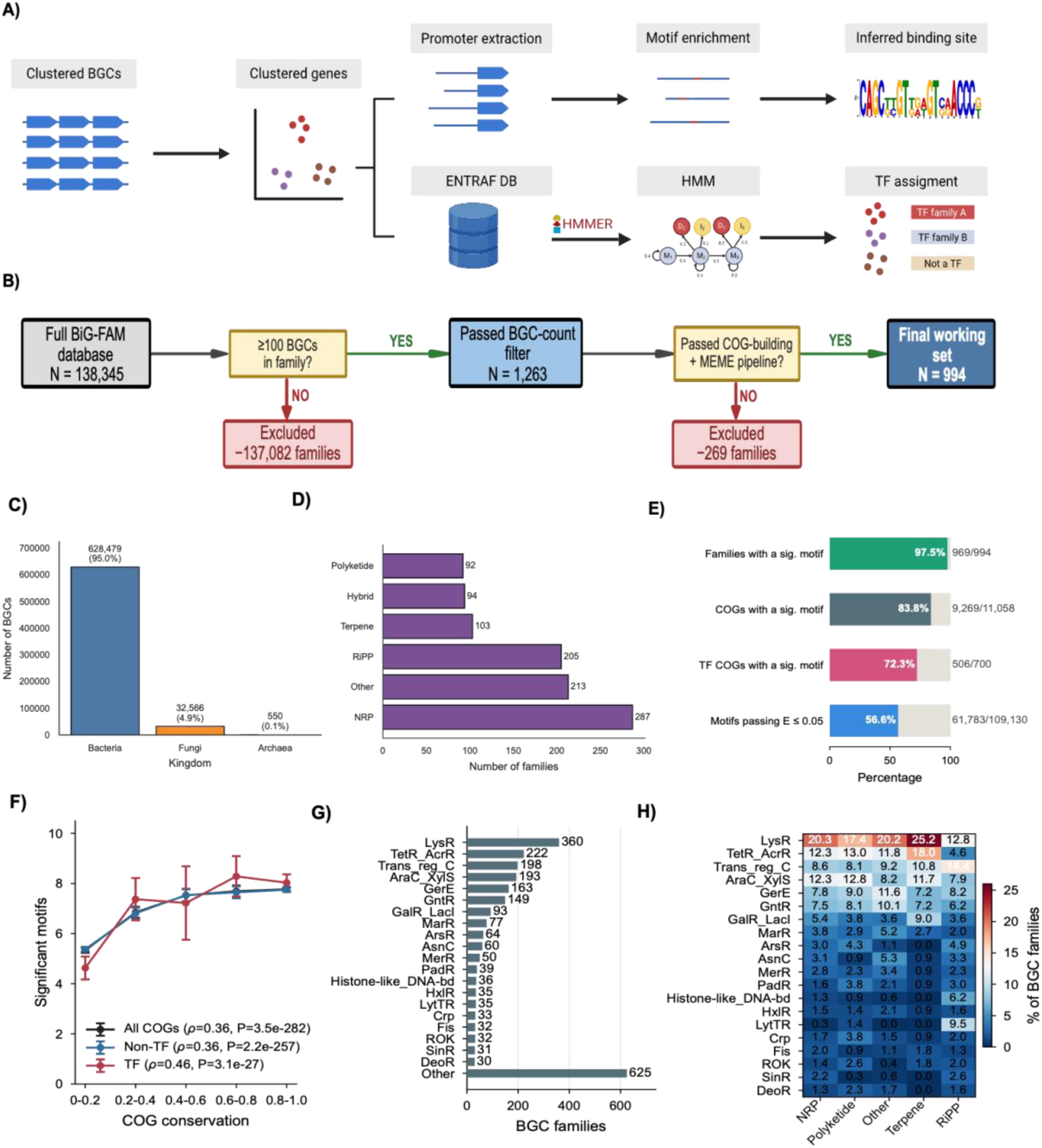
Pipeline overview, dataset composition and the pathway-specific transcription factor landscape. **(A)** Schematic of the analysis pipeline. Member BGCs of each gene cluster family are partitioned into COGs, upstream promoter regions are extracted, and *de novo* motif discovery yields inferred transcription factor binding sites (upper arm). In parallel, HMM profiles built from the ENTRAF database are searched against all BGC-encoded genes to assign each to a TF family where matched, enabling predicted sites to be linked to candidate regulators (lower arm). **(B**) Data flow and family counts from the full BiG-FAM database through the ≥100-BGC size filter to the final analysed set. **(C)** Distribution of BGCs by source kingdom across the analysed families, with counts and percentages. **(D)** Distribution of dominant BGC structural classes across the analysed families. **(E)** Coverage of the pipeline output under the zoops_maxw30 setting, showing the proportion of families, COGs and TF-encoding COGs with at least one significant motif, and the fraction of all discovered motifs passing significance. Significant motifs are those with E ≤ 0.05. **(F)** Relationship between COG conservation and the number of significant motifs recovered (maximum 10 per run), shown for all COGs, non-TF COGs and TF-encoding COGs. Points and error bars show mean ± s.d. per bin; Spearman correlation coefficients (ρ) and P-values (P) are inset. **(G)** Number of BGC families assigned to each TF family, ranked by family count; families outside the top 20 are grouped as ‘Other’. **(H)** Column-normalised representation of BGC families per TF family (top 20) stratified by BGC class; families with hybrid class designations contribute to each constituent class. **Alt text:** Multi-panel figure. **(A)** Flow diagram of the pipeline: BGC family members are split into orthologous gene groups, upstream regions are extracted and searched for motifs, while a parallel arm assigns transcription factor families using HMM profiles. **(B)** Flow chart of family counts from the full BiG-FAM database through the size filter to the 994 analysed families. **(C, D)** Distributions of analysed BGCs by kingdom and by biosynthetic class. **(E)** Proportions of families, COGs and TF-encoding COGs with significant motifs. **(F)** Line plots showing the number of significant motifs rising with COG conservation, more steeply for TF COGs. **(G)** Bar chart of BGC families per TF family, led by LysR. **(H)** Heatmap of TF family representation across biosynthetic classes.

Of the 138,345 GCFs in BiG-FAM, 58.3% contain a single BGC and 88.1% contain five or fewer. Of the 1,263 families with at least 100 members, 994 contained at least one COG with four or more dereplicated upstream sequences and were analysed (**Figure 1B**).

The six MEME parameter combinations recovered complementary but non-identical sets of motifs, with no single setting dominating (**Supplementary Figure S1**). ZOOPS with a maximum width of 30 bp was adopted as the primary setting because ZOOPS corresponds to canonical single-site binding and a 30-bp ceiling allows MEME to select the optimal width up to the length of the longest characterised bacterial operators. Motifs from all six settings are available for download.

The analysed families comprise 628,479 bacterial (95.0%), 32,566 fungal (4.9%) and 550 archaeal (0.1%) BGCs (Figure 1C), similar to the full database (**Supplementary Figure S2A**). Non-ribosomal peptide (NRP) families are the most numerous (287, 28.9%), followed by ‘Other’ (213, 21.4%), RiPP (205, 20.6%), terpene (103, 10.4%), hybrid (94, 9.5%) and polyketide (92, 9.3%) families (**Figure 1D**). Polyketide families are depleted relative to BiG-FAM (18.4% of families), most likely because the variable architecture of polyketide synthase clusters fragments them into many small GCFs, and saccharide and alkaloid families, which are rare in BiG-FAM, are absent (**Supplementary Figure S2B**).

### Pipeline output

Across the 994 families, 61,783 of 109,130 motifs discovered under the primary setting (56.6%) passed the significance threshold of E ≤ 0.05 (**Figure 1E**). At least one significant motif was recovered in 969 of 994 families (97.5%) and in 9,269 of 11,058 COGs (83.8%). Among COGs encoding a predicted TF, 506 of 700 (72.3%) carried a significant upstream motif. Most families comprise a modest number of COGs, consistent with compact pathways, while a smaller subset contains many more, reflecting larger and more architecturally complex clusters (**Supplementary Figure S3**).

We examined how motif recovery relates to COG conservation, and whether this differs between TF-encoding and other COGs (**Figure 1F**). In both groups the number of significant motifs increased with the fraction of the family’s BGCs in which the COG was present, as expected, since widely conserved COGs contribute more upstream sequences and therefore more power for motif detection. The relationship was stronger for TF COGs (Spearman ρ = 0.46) than for non-TF COGs (ρ = 0.36; P < 10⁻¹⁵ for both), although the difference is modest. Gene organisation offers a likely explanation. TF genes were about half as likely as other cluster genes to be operon-internal (same-strand upstream neighbour within 50 bp: 7.1% vs 22.8% of genes; 9–17% vs 20–26% of genes per COG across COG-size quintiles), and among the larger, more conserved COGs, TF genes had longer upstream intergenic regions than other genes (**Supplementary Figure S4**). Conserved regulators therefore tend to carry their own promoter regions, in which conserved sites can be detected, whereas roughly one in four biosynthetic genes lies within an operon and its upstream window contains mostly coding sequence. The steeper relationship for TF COGs reflects where promoters are located rather than how strongly regulators are regulated.

### Landscape of pathway-specific transcription factors across BGC families

We next characterised the regulators encoded within the analysed families. HMM-based annotation across the 994 GCFs identified 2,986 high-confidence TF COGs spanning 117 TF families, and a dominant PSTF was assigned to 921 GCFs (see Materials and Methods).

LysR-family regulators were the most prevalent PSTFs, dominant in 360 GCFs encompassing 131,441 BGCs, the highest by both measures (**Figure 1G**). TetR/AcrR, the Trans_reg_C two-component response regulators and AraC/XylS followed by family count, with GerE and GntR also prominent.

Normalising within each BGC class revealed several class-specific patterns (**Figure 1H**). LysR was the most prevalent PSTF in every class (12.8–25.2% of families), consistent with its role as a generalist regulator. Against this background, several families were strongly class restricted. LytTR was almost exclusive to RiPP families (9.5%, below 1.5% in all other classes), in line with its established role in bacteriocin regulation (34, 35), and Histone-like DNA-binding proteins were likewise enriched in RiPP (6.2%). Two-component response regulators (Trans_reg_C) leaned modestly towards RiPP (14.4%) and terpene (10.8%) over NRP (8.6%) and polyketide (8.1%) families.

Terpene and RiPP families showed the most distinctive profiles. Terpenes were dominated by two TF families, LysR (25.2%) and TetR/AcrR (18.0%), with several others absent altogether (ArsR, LytTR, Histone-like, SinR and DeoR), indicating a comparatively narrow regulatory repertoire. RiPPs were the only class in which LytTR, Histone-like proteins and Trans_reg_C were all well represented. Although the percentage differences between classes are modest for most TF families, these patterns indicate that the regulatory repertoire is not uniform across biosynthetic classes. Some classes rely mainly on a small set of generalist regulators, while others recruit more class-specific ones.

### Recovery of experimentally characterised binding sites

To test whether de novo motifs correspond to known regulatory elements, we examined GCFs containing MIBiG-curated BGCs with experimentally characterised operators, covering ectoine and siderophore clusters (**Table 1**, **Figure 2**, **Supplementary Figures S5–S7**). Each reference operator was compared by TOMTOM against all motifs predicted for the family and was not used at any stage of discovery.

**Figure 2:**
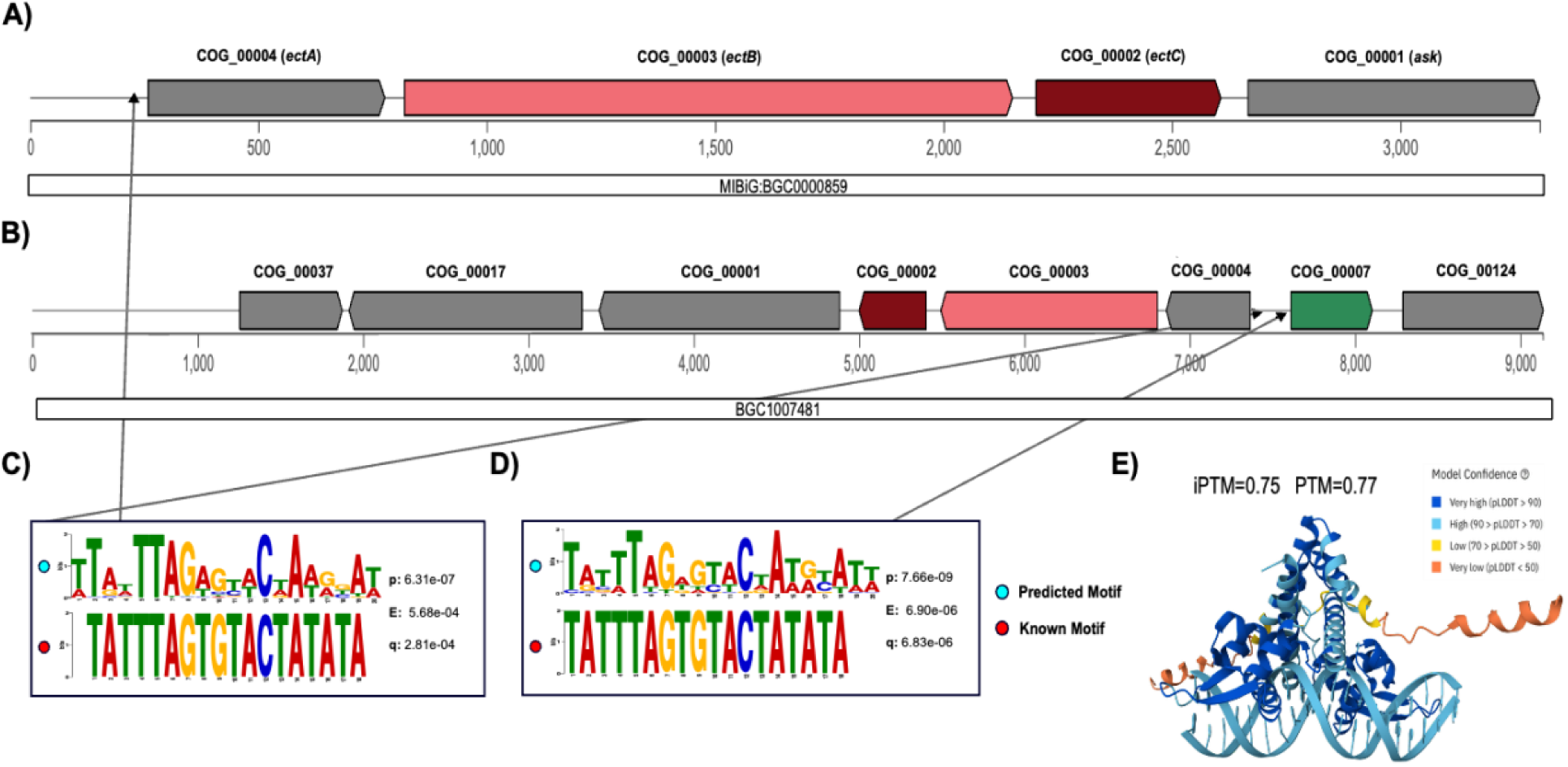
Recovery of the EctR1 operator in ectoine BGC families. **(A)** Genomic architecture of the MIBiG anchor BGC0000859 (*Methylomicrobium alcaliphilum*, FAM9025): COG_00004/*ectA*, COG_00003/*ectB*, COG_00002/*ectC* and COG_00001/*ask*. **(B)** Genomic architecture of BGC1007481 (*Teredinibacter sp.,* FAM9025); the four biosynthetic COGs are present alongside the MarR-family regulator COG_00007 and additional COGs absent from the anchor. Genes in (A) and (B) are coloured by predicted function (burgundy, biosynthetic; pink, biosynthetic-additional; green, regulatory; grey, other). **(C)** TOMTOM comparison of motif_1 upstream of COG_00004 (*ectA*) in FAM9025 (blue, pipeline-predicted) against the experimentally characterised EctR1 binding site TATTTAGTGTACTATATA (red); p, p-value; E, E-value; q, q-value. **(D)** TOMTOM comparison of motif_5 upstream of the MarR regulator COG_00007 against the EctR1 site. **(E)** AlphaFold3 model of the COG_00007 MarR homodimer in complex with the *ectA* motif (COG_00004 motif_1), coloured by pLDDT (ipTM = 0.75, pTM = 0.77). The scrambled-control complex scored identically (ipTM = 0.75); the monomer–DNA model distinguishes real from scrambled motif (ipTM 0.47 vs 0.10; Supplementary Table S2). **Alt text:** Multi-panel figure on ectoine clusters. **(A, B)** Gene arrow diagrams of two ectoine clusters from FAM9025, coloured by gene function; the *Teredinibacter* cluster in **(B)** carries an additional MarR-family regulator gene. **(C, D)** Paired sequence logos comparing pipeline-predicted motifs upstream of ectA and of the MarR regulator with the experimentally characterised EctR1 binding site, with matching statistics. **(E)** AlphaFold3 structural model of the MarR homodimer bound to DNA carrying the *ectA* motif, coloured by prediction confidence.

**Table 1:**
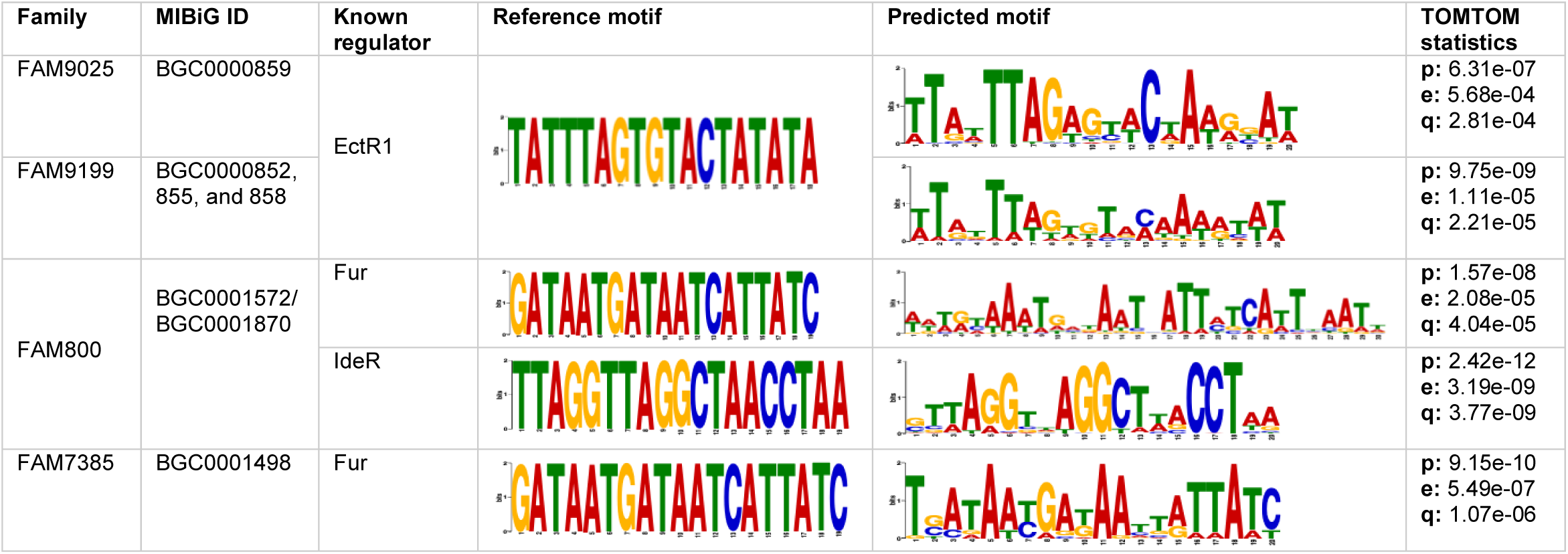
Recovery of experimentally characterised TFBSs by the pipeline. Comparison of experimentally characterised TFBSs with motifs independently recovered by the pipeline. For each BGC family, the MIBiG-curated anchor cluster and product and the experimentally validated regulator are listed alongside the reference binding site used as the TOMTOM query. TOMTOM values report the significance of the match between the pipeline-predicted motif and the reference site (p, raw p-value; e, E-value; q, multiple-testing-corrected q-value).

| Family | MIBiG ID | Known regulator | Reference motif | Predicted motif | TOMTOM statistics |
|---|---|---|---|---|---|
| FAM9025 | BGC0000859 | EctR1 |  |  | **p:** 6.31e-07<br>**e:** 5.68e-04<br>**q:** 2.81e-04 |
| FAM9199 | BGC0000852, 855, and 858 |  |  |  | **p:** 9.75e-09<br>**e:** 1.11e-05<br>**q:** 2.21e-05 |
| FAM800 | BGC0001572/<br>BGC0001870 | Fur |  |  | **p:** 1.57e-08<br>**e:** 2.08e-05<br>**q:** 4.04e-05 |
|  |  | IdeR |  |  | **p:** 2.42e-12<br>**e:** 3.19e-09<br>**q:** 3.77e-09 |
| FAM7385 | BGC0001498 | Fur |  |  | **p:** 9.15e-10<br>**e:** 5.49e-07<br>**q:** 1.07e-06 |

Ectoine biosynthetic genes are typically organised as an osmolarity-induced *ectABC* operon (36). Three regulators have been characterised. The MarR-family repressor EctR1, which binds a pseudopalindromic operator overlapping an *ectABC* promoter (28), the related repressor CosR, whose binding site is unresolved (37), and the global nitrogen regulator GlnR, which binds the actinomycete consensus gTnAc-n₆-GaAAc-n₆ (38–40).

The EctR1 operator was recovered upstream of *ectA* (COG_00004) in two families. FAM9025 contains the *Methylomicrobium alcaliphilum* cluster in which EctR1 was characterised [MIBiG BGC0000859] (q = 2.8×10⁻⁴, **Figure 2A, C**). FAM9199 comprises ectoine clusters from other species [MIBiG BGC0000852, BGC0000855, BGC0000858] in which the operator has not been experimentally defined, and a matching motif was again recovered upstream of *ectA* (q = 2.2×10⁻⁵, **Supplementary Figure S5**). No EctR1 match was found upstream of *ectB*, *ectC* or *ask* in either family, consistent with a single operator controlling the co-transcribed unit (28, 36).

In FAM9025, BGC1007481 (*Teredinibacter* sp.) additionally encodes a MarR-family regulator (COG_00007) with 70.5% amino-acid identity to EctR1 (**Figure 2B**), and an EctR1-like motif was recovered upstream of it (q = 6.8×10⁻⁶, **Figure 2D**), consistent with autoregulation, a common feature of MarR-family repressors (41–43).

AlphaFold3 modelling of this regulator (**Supplementary Table S2**, **Supplementary Figure S8**) gave a confident MarR fold and dimer (monomer pTM = 0.80, dimer ipTM = 0.76). The dimer–*ectA* operator complex was modelled confidently (ipTM = 0.75, pTM = 0.77, **Figure 2E**) but scored identically against a composition-matched scrambled control, because the protein–protein interface dominates the dimer score. The monomer–DNA model separated real from scrambled DNA (ipTM 0.47 vs 0.10). In further cases in uncharacterised clusters, AlphaFold3 frequently failed to distinguish real from scrambled sequences (**Supplementary Tables S3–S6**, **Supplementary Figures S9–S10**).

FAM238, which contains the *Streptomyces anulatus* hydroxyectoine cluster [MIBiG BGC0000853], yielded strongly conserved motifs upstream of all four genes (E < 10⁻¹¹⁶, **Supplementary Figure S6**), none matching the EctR1 operator or the GlnR consensus, suggesting an uncharacterised regulatory architecture.

Siderophore biosynthesis is repressed under iron-replete conditions by the ferric uptake regulator (Fur) and analogous regulators such as IdeR (44–47). Predicted motifs were compared against the *Escherichia coli* Fur box (GATAATGATAATCATTATC) and the *Mycobacterium tuberculosis* IdeR operator (TTAGGTTAGGCTAACCTAA) (48–51).

FAM800 groups desferrioxamine-type clusters from Proteobacteria, Bacteroidota and Actinobacteria, including two MIBiG clusters [BGC0001572, BGC0001870]. Two distinct motifs were recovered upstream of dfoJ (COG_00001), one reproducing the IdeR box (q = 3.8×10⁻⁹) and one containing the Fur box (q = 4.0×10⁻⁵), each under at least three of the six MEME settings (**Table 1**, **Supplementary Figure S7A–B**). The iron box occurred in nearly all actinobacterial members and in no proteobacterial member, whereas the Fur box occurred in most proteobacterial members, including both MIBiG anchors, and in *Elizabethkingia*, but in no actinobacterial member. No sequence carried both. In the purely proteobacterial FAM7385 [MIBiG BGC0001498], a Fur box was recovered upstream of iucA (COG_00005, q = 1.1×10⁻⁶) with no IdeR match (**Supplementary Figure S7D–E**).

Across the four families with characterised operators, the known site was recovered upstream of the expected target gene and not upstream of co-transcribed genes.

### Sequence similarity network analysis

SSNs combining pipeline-annotated TF sequences with PRODORIC-characterised regulators were built for the 20 most prevalent PSTF families (**Figure 3A**, Methods). Four are presented here, and all 20 are available in the resource.

**Figure 3:**
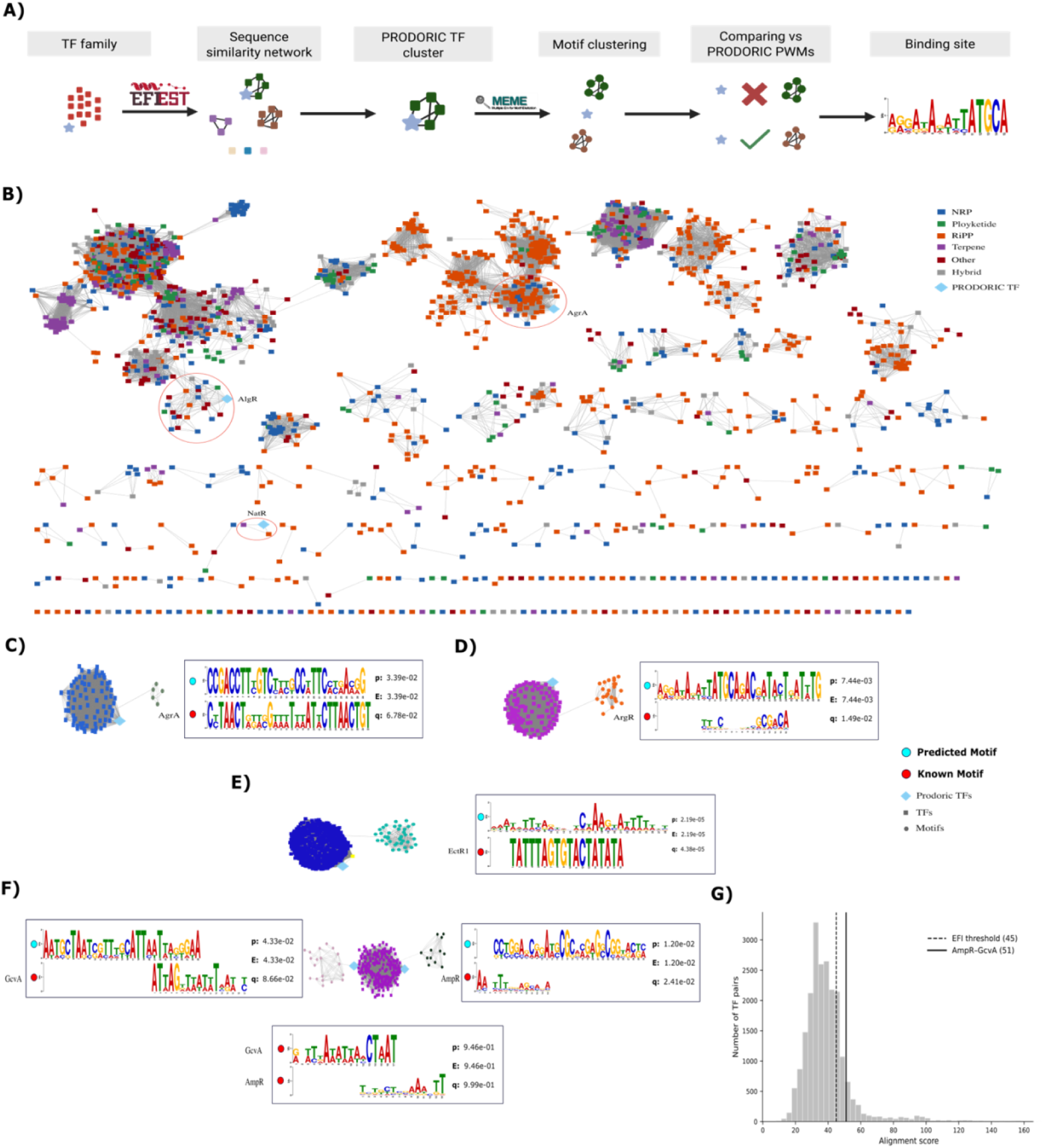
Sequence similarity network analysis of LytTR, AraC/XylS, MarR and LysR transcription factor families. **(A)** Schematic of the analysis. TF sequences of a family are combined with experimentally characterised regulators from PRODORIC and clustered into an SSN. Within each SSN TF cluster, upstream motifs from the member BGC families are clustered by similarity, and conserved motif clusters are compared against the PRODORIC binding-site PWM of any characterised regulator in the same SSN TF cluster. **(B)** SSN of all LytTR-annotated TF sequences (alignment-score threshold 40). Nodes are TF sequences coloured by the BGC class of the source BGC (blue, NRP; green, polyketide; orange, RiPP; purple, terpene; red, other; grey, hybrid); light-blue diamonds are PRODORIC TFs. Circles mark the SSN TF clusters containing AgrA, AlgR and NatR. **(C–F)** Combined TF and motif networks for individual SSN TF clusters. TF nodes (squares), motif nodes (circles) and PRODORIC TFs are light-blue diamonds. Each is paired with the TOMTOM comparison of the cluster’s best-matching conserved motif (blue circle, pipeline-predicted) against the experimentally characterised binding site (red circle; p, p-value; E, E-value; q, q-value). (C) LytTR SSN TF cluster 3, containing AgrA (q = 0.068). (D) AraC/XylS SSN TF cluster 10, containing ArgR (q = 0.015). (E) MarR SSN TF cluster 0, containing *M. alcaliphilum* EctR1 and the *Teredinibacter sp*. COG_00007 regulator of FAM9025 (yellow square); the cluster’s conserved motif matches the EctR1 binding site (q = 4.4×10⁻⁵). (F) LysR SSN TF cluster 3, containing GcvA (left) and AmpR (right), each matched by a separate conserved motif cluster (q = 0.087 and 0.024); centre, direct TOMTOM comparison of the GcvA and AmpR binding sites (q = 0.999). **(G)** Distribution of pairwise alignment scores among TF pairs in LysR SSN TF cluster 3; dashed line, SSN construction threshold (45); solid line, AmpR– GcvA score (51). **Alt text:** Multi-panel figure of transcription factor networks. **(A)** Schematic of the method linking TF sequence similarity networks to clustered upstream motifs and characterised binding sites. **(B)** Network of LytTR family sequences coloured by BGC class, dominated by RiPP clusters, with characterised regulators marked. **(C–F)** Networks combining TF nodes and motif nodes for individual TF subfamilies, each paired with sequence logos comparing the best conserved motif with a characterised binding site: **(C)** AgrA, **(D)** ArgR, **(E)** EctR1, and **(F)** the LysR subfamily containing AmpR and GcvA with two separate motif matches. **(G)** Histogram of pairwise alignment scores within the LysR subfamily, marking the network threshold and the AmpR–GcvA score.

The LytTR network resolved into 133 SSN TF clusters drawn predominantly from RiPP BGCs (**Figure 3B**), consistent with **Figure 1H**, and included AgrA (MX000357), AlgR (MX000101) and NatR (MX000197). SSN TF cluster 3, containing the *Staphylococcus aureus* quorum-sensing regulator AgrA, gave a borderline match to the AgrA binding site (q = 0.068, **Figure 3C**). AgrA is a global rather than pathway-specific regulator, and BGC-associated sites may diverge from those of core regulons (10, 52).

The AraC/XylS network resolved into 1,006 multi-member SSN TF clusters and included 15 PRODORIC TFs (**Supplementary Figure S11**). SSN TF cluster 10, enriched for TFs from NRP BGCs, contained the arginine-responsive *Pseudomonas aeruginosa* regulator ArgR (MX000007) (53), and its conserved motif cluster 29 (family coverage 1.0) matched the ArgR binding site (q = 0.015, **Figure 3D**).

The MarR network resolved into 645 SSN TF clusters and included eight PRODORIC TFs plus EctR1 with its literature-derived binding site (**Supplementary Figure S12**) (29). SSN TF cluster 0 contained EctR1 and the *Teredinibacter* sp. regulator of FAM9025, and its conserved motif cluster matched the EctR1 binding site (q = 4.4×10⁻⁵, **Figure 3E**), recovering the same operator independently of the family-level analysis.

The LysR network resolved into 890 SSN TF clusters spanning all major BGC classes and included 19 PRODORIC TFs (**Supplementary Figure S13**). SSN TF cluster 3 contained AmpR (MX000276), which controls β-lactamase expression in *Pseudomonas*, and GcvA (MX000143), which regulates the glycine cleavage system in *E. coli* (54, 55). The two proteins fall in the same SSN TF cluster but recognise unrelated DNA sequences (TOMTOM q = 0.999 between their binding sites). The pipeline recovered two separate conserved motif clusters within this subfamily without prior knowledge, one matching the AmpR site (q = 0.024) and the other showing a borderline match with the GcvA site (q = 0.087) (**Figure 3F**). Sequence similarity between regulators therefore does not imply a shared binding site (pairwise alignment score 51, above the network threshold of 45, **Figure 3G**), and the motif network can retain more than one conserved motif within a single TF subfamily rather than collapsing them into one consensus.

Across the 20 networks, 779 SSN TF clusters carry at least one conserved motif cluster, providing a prioritised catalogue of TF subfamilies with candidate pathway-specific binding sites (**Supplementary Figures S14–S17**).

### The BGC-TFBS resource

All results are available through the BGC-TFBS web resource (https://bgc-tfbs.sites.er.kcl.ac.uk/), which can be explored from either the BGC or TF perspective (**Figure 4** BGC families can be searched by BiG-FAM identifier, MIBiG accession, assembly accession or taxon, and filtered by product class, family size and MIBiG membership (**Figure 4A**). Each family page summarises its composition, pathway-specific TFs and MIBiG reference clusters, and displays member BGCs as COG-coloured gene tracks. Selecting a COG reveals its predicted motifs ranked by MEME E-value, with sequence logos, consensus sequences and the number of dereplicated upstream sequences containing each motif (**Figure 4B**). The 20 PSTF families can be browsed by SSN TF cluster, showing taxonomic and BGC-class composition, conserved motif clusters with family coverage and representative logos, and matches to PRODORIC-characterised binding sites (**Figure 4C**). The underlying tables, motifs from all six MEME settings and network files are also available for download.

**Figure 4:**
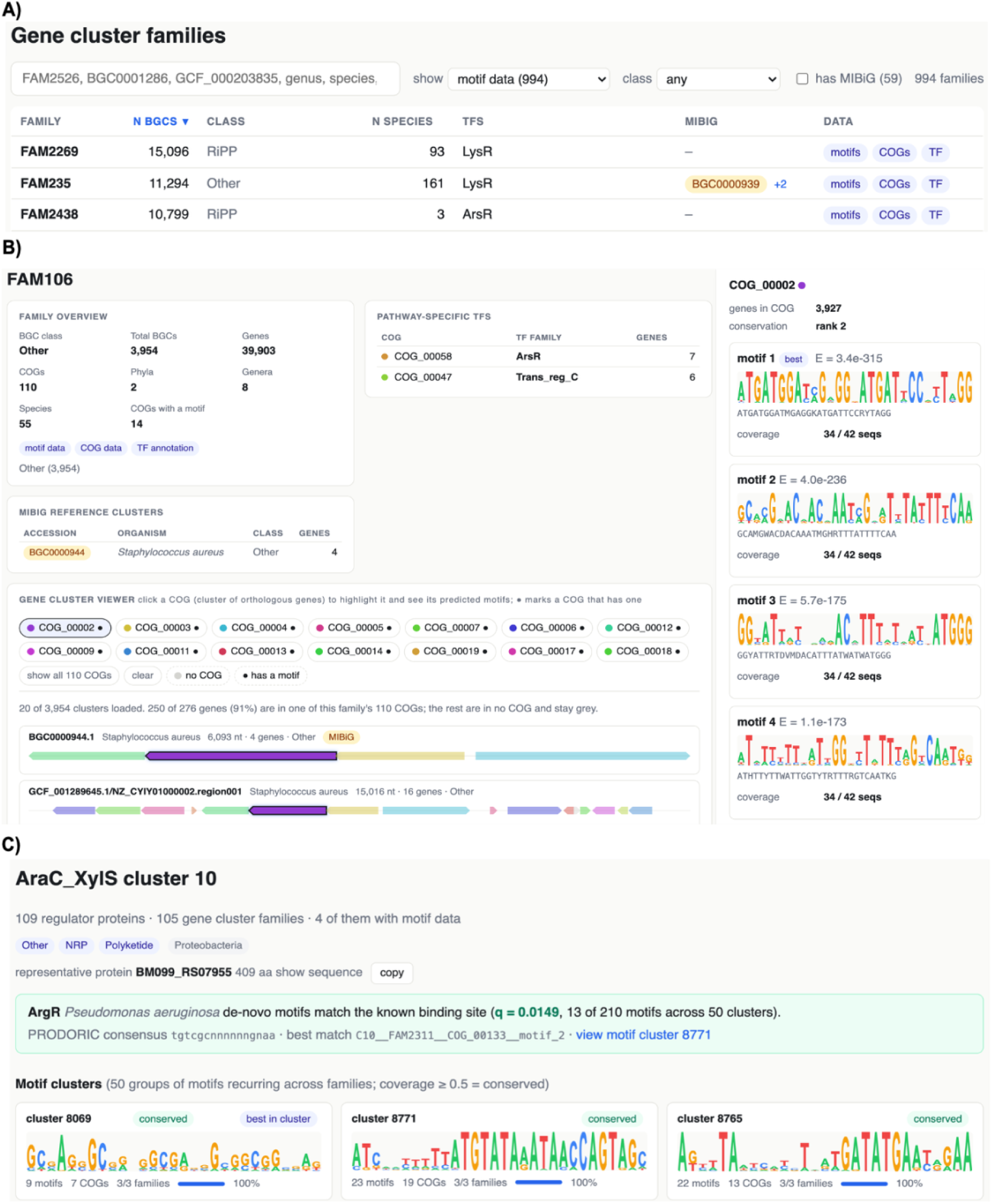
The BGC-TFBS web resource. **(A)** Families can be searched by BiG-FAM identifier, MIBiG accession, assembly accession or taxon and filtered by product class and MIBiG membership. Each row gives family size, dominant BGC class, species count, dominant PSTF family and MIBiG members. **(B)** Family page for FAM106 (3,954 BGCs), showing the family overview, its pathway-specific TFs (ArsR, Trans_reg_C), the MIBiG reference cluster BGC0000944, and the gene cluster viewer with genes coloured by COG. Selecting a COG_00002 opens its predicted motifs ranked by MEME E-value, each with sequence logo, consensus and the number of dereplicated upstream sequences carrying it (34 of 42). **(C)** Regulator view for AraC/XylS SSN TF cluster 10, showing the cluster’s composition, its match to the PRODORIC-characterised ArgR binding site (q = 0.0149, Figure 3D) and its conserved motif clusters (family coverage ≥ 0.5) with logos, member counts and coverage. All tables, motifs from all six MEME settings and network files are available from the Downloads page. **Alt text:** Three screenshots of the BGC-TFBS website. **(A)** Searchable table of BGC families. **(B)** Family page showing COG-coloured gene clusters with predicted motif logos. **(C)** Regulator page for an AraC/XylS subfamily showing its conserved motifs and match to the ArgR binding site.

## DISCUSSION

We have built a comparative-genomic pipeline for predicting candidate TFBSs across GCFs and applied it to 994 families from BiG-FAM. Because motif discovery is performed on the upstream regions of orthologous genes, candidate binding sites are found without prior knowledge of the regulator, and HMM-based annotation of cluster-encoded proteins links them to candidate TF families.

To our knowledge, the distribution of PSTF families across BGC classes has not previously been described at GCF scale. LysR-type regulators occurred in every class, in line with their known versatility (56–59), whereas others were class-restricted, most clearly LytTR in RiPP clusters (34, 35). Most between-class differences are modest, and we interpret them as associations between regulator families and pathway types rather than evidence of co-evolution.

The recovery tests provide the strongest evidence that predicted motifs correspond to regulatory elements. The EctR1 operator was recovered upstream of ectA both in FAM9025, which contains the cluster in which EctR1 was characterised, and in FAM9199, whose members come from species in which the operator has not been experimentally defined (28). The pipeline therefore detects conserved regulatory architecture rather than re-finding a reference sequence in its source genome. In FAM800, the pipeline recovered two mutually exclusive motifs upstream of the same orthologous gene, a Fur box in Proteobacteria and Bacteroidota and a DtxR-family iron box in Actinobacteria, each restricted to the lineage in which its cognate regulator operates (60–65). An orthologous group can therefore carry more than one regulatory solution for the same biosynthetic step. De novo discovery separates them, whereas scanning against a single reference matrix would recover only one.

The paired TF and motif networks ask a different question, which is how regulator sequence relates to operator specificity. Clustering TF sequences into subfamilies and, separately, clustering the upstream motifs of their member families lets the two be compared directly. In a LysR subfamily containing AmpR and GcvA, the two proteins are similar enough to co-cluster but recognise unrelated sequences, and the pipeline recovered a distinct conserved motif cluster matching each of them without knowing either in advance. What the networks add is that motif clustering can hold both specificities side by side instead of collapsing a subfamily into one consensus. In the MarR network, the subfamily containing EctR1 and the *Teredinibacter* regulator from FAM9025 produced a conserved motif matching the EctR1 operator, so the same site was reached both from inside one family and from across the BGC families of an entire TF subfamily. The AraC/XylS network gave a further match, with the subfamily containing the arginine-responsive regulator ArgR of *Pseudomonas aeruginosa* carrying a conserved motif that matched the ArgR operator across all of its member families (53).

The pipeline fills a gap left by existing tools. The antiSMASH TFBS Finder scans an individual cluster against known binding-site models (15), and COMMBAT does so at scale, combining matches to more than 4,000 TF matrices with genomic context and gene function across over 400,000 BGCs to predict which TFs act on which clusters (16, 66). Both give the identity of the regulator directly, but only when a matrix for it exists, and many BGC-associated regulators have none. *De novo* discovery removes that dependence but needs a set of co-regulated promoters as input, which the AURTHO framework obtained for TF-encoding genes by comparing the upstream regions of orthologous regulators across species (18). We apply the same logic to every gene in a cluster, with orthologues within a GCF supplying the co-regulated set. The regulator is then assigned not from a matrix but from conservation, by annotating cluster-encoded TFs and asking which motifs recur across the families of a TF subfamily, and the recovery cases show that this reaches the right operator where one is known.

Structural modelling with AlphaFold3 was of limited use for establishing specificity. Monomer interface scores separated the EctR1 operator from a composition-matched control, but in other cases the method failed to distinguish predicted motifs from scrambled sequences, and dimer scores were dominated by the protein–protein interface. Structural prediction is therefore best treated as corroborating evidence for strong sequence-level predictions rather than as a test of binding specificity.

The main limitation is the requirement of at least 100 BGCs per GCF. The threshold is a proxy for sequence diversity: MEME needs several distinct upstream sequences per COG, and dereplication at 90% identity removes most of a family’s members, so a lower cutoff would mainly admit families that fail at motif discovery. As a result, polyketide families are under-represented and saccharide and alkaloid families are absent, so the catalogue is not a comprehensive view of BGC regulation. Within families, sites can only be found inside the extracted upstream window. Around one in four non-regulatory genes is operon-internal, so its window is mostly coding sequence, and distal or intragenic sites are out of reach. TF annotation rests on DNA-binding domain profiles, which identify a cluster-encoded regulator as a candidate PSTF but do not show that it regulates its own cluster. Because these profiles derive from bacterial regulators, fungal clusters receive motifs but no regulator assignment. Finally, a conserved motif matching a characterised operator is strong evidence for a regulatory element but not proof of binding.

The value of a prioritised catalogue is that it reduces a large search space to a testable set. Predicted motifs can be tested by EMSA, ChIP or DAP-seq (67, 68), and combined with transcriptomic data to test whether predicted regulons behave as regulons. By mapping candidate binding sites across BGC families alongside their candidate regulators, this resource provides a starting point for the experimental dissection of BGC regulation and the activation of silent clusters.

## ACKNOWLEDGEMENTS

The authors acknowledge use of the research computing facility at King’s College London, King’s Computational Research, Engineering and Technology Environment (CREATE) (69).

## AUTHOR CONTRIBUTIONS

The concept for this study was developed by I.M., F.G.G. and S.S. I.M. designed and implemented the pipeline, performed all analyses, built the web resource, prepared the figures and wrote the first draft of the manuscript. F.G.G. contributed significantly to the design of the analyses and the interpretation of the results. F.C. contributed to editing the manuscript. S.S. and M.G. contributed to conceptualisation, secured funding, supervised the project and assisted with writing and editing. All authors read and approved the final version of the manuscript.

## DATA AVAILABILITY

The data underlying this article are available in Zenodo at https://doi.org/10.5281/zenodo.22766286. The motif discovery pipeline is available at https://github.com/sysbiomelab/Motif-Miner and archived at https://doi.org/10.5281/zenodo.23067090. The code for the BGC-TFBS web resource is available at https://github.com/sysbiomelab/BGC-TFBS and archived at https://doi.org/10.5281/zenodo.23066946. The resource itself is freely accessible at https://bgc-tfbs.sites.er.kcl.ac.uk/ without registration.

## KEY POINTS

- Orthologous genes within biosynthetic gene cluster families provide co-regulated upstream regions for de novo discovery of transcription factor binding sites without prior knowledge of the regulator.
- Applied to 994 BiG-FAM families, the pipeline recovered experimentally characterised operators.
- LysR-type regulators are the most prevalent pathway-specific transcription factors in every biosynthetic class, whereas families such as LytTR are largely restricted to RiPP clusters.
- Combining transcription factor sequence similarity networks with motif clustering can resolve more than one conserved motif within a single regulator subfamily.

