## Supplemental file for "Transcription Factor Binding Site Prediction in Microbial Biosynthetic Gene Cluster Families"

### Supplementary Figure:

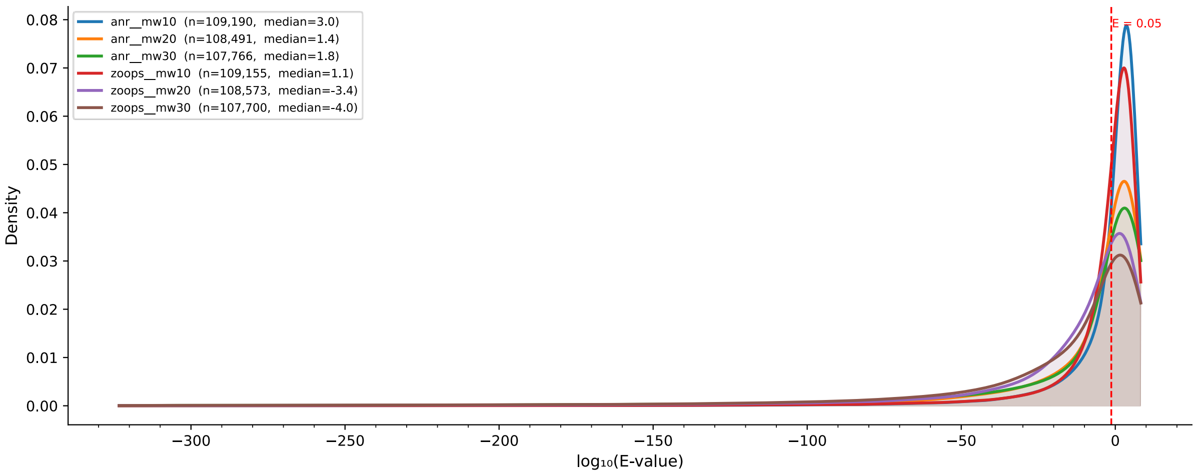


**Figure S1: Comparison of MEME motif E-value distributions across parameter settings.** Kernel density estimates of $\log_{10}$-transformed motif E-values are shown for each MEME parameter combination, with shaded areas representing the estimated probability density. The legend indicates the number of motifs (n) and median $\log_{10}$(E-value) for each parameter setting. A vertical dashed red line marks the significance threshold of E = 0.05 ($\log_{10}$ = −1.30)

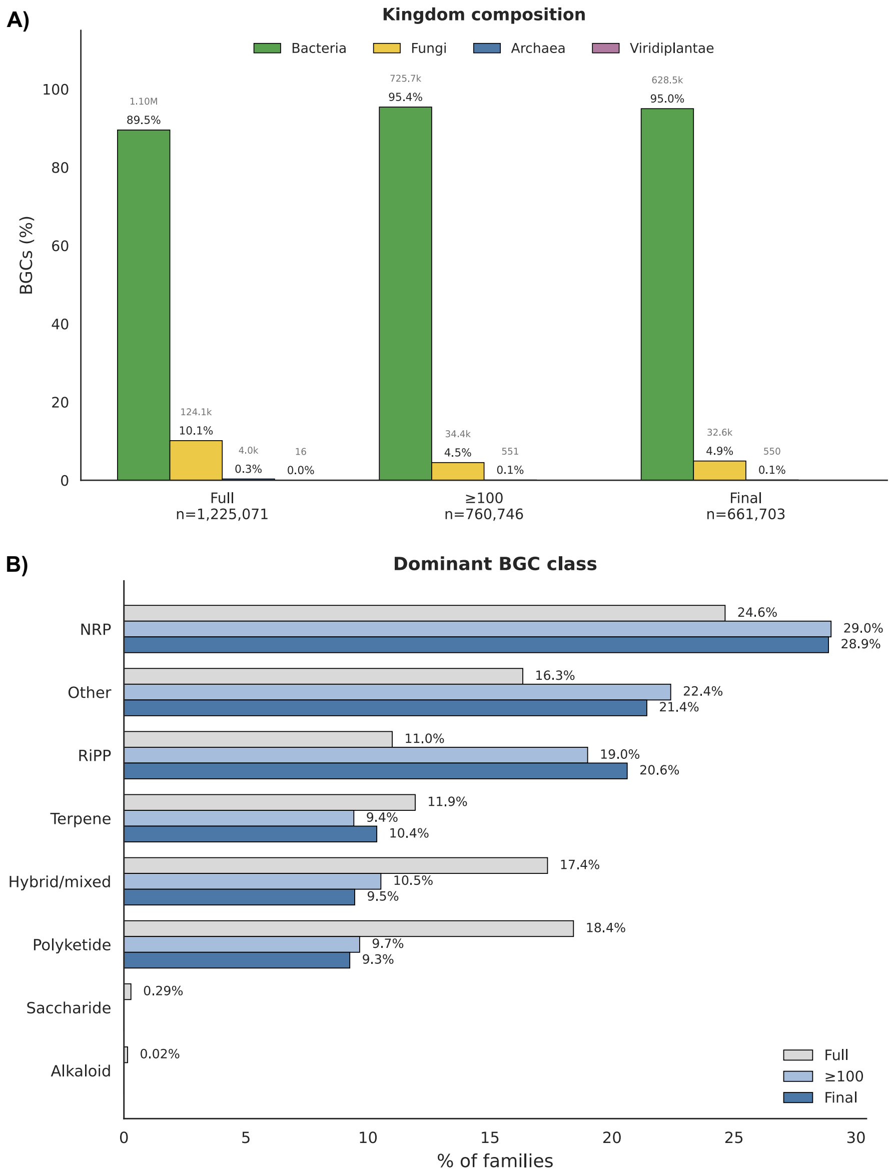


**Figure S2: Taxonomic and biosynthetic class composition of BGCs through the filtering steps. (A)** Kingdom composition of BGCs at three stages: the full BiG-FAM database (138,345 families), families with ≥100 member BGCs (1,263), and the analysed set with motif output (994). **(B)** Distribution of dominant BGC classes across families at the same three stages.

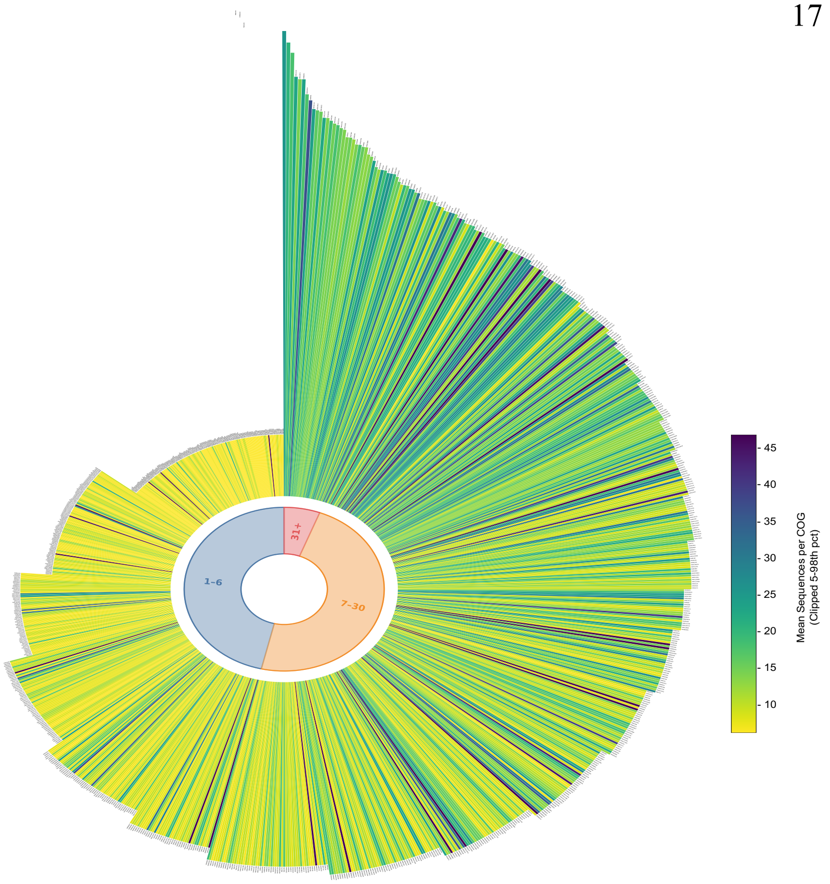


**Figure S3: Number of COGs per gene cluster family.** Circular bar plot of all 994 gene cluster families (GCFs), each represented by a single radial bar. Bar height denotes the number of COGs per family. Bar colour indicates the mean number of upstream sequences per COG (clipped to the 5th–98th percentile). The inner ring partitions families into three COG-complexity bands (1–6, 7–30, and 31+ COGs).

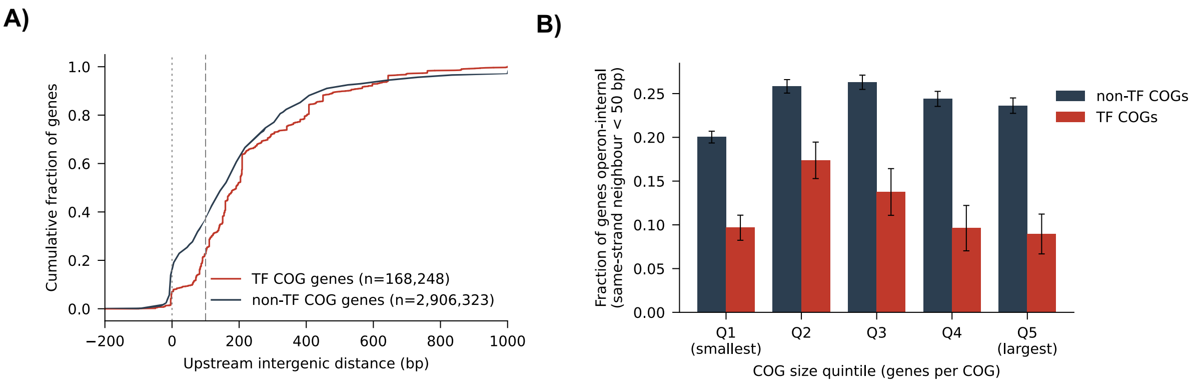


**Figure S4: Upstream intergenic distance of TF-encoding and other cluster genes. (A)** Cumulative distribution of upstream intergenic distance for genes in TF COGs (n = 168,248) and non-TF COGs (n = 2,906,323) across the 994 analysed families. Upstream distance is measured from the gene's start codon to the end of the nearest preceding gene in the BGC (negative values indicate overlapping genes); genes at BGC boundaries with no upstream neighbour (n = 212,493) are excluded. Dotted line, 0 bp; dashed line, 100 bp, the minimum upstream window used for motif discovery. Median distance 191 bp (TF) vs 150 bp (non-TF); Mann–Whitney P < $10^{-15}$. **(B)** Mean fraction of genes per COG classed as operon-internal (same-strand upstream neighbour within 50 bp) by COG-size quintile, for TF and non-TF COGs.

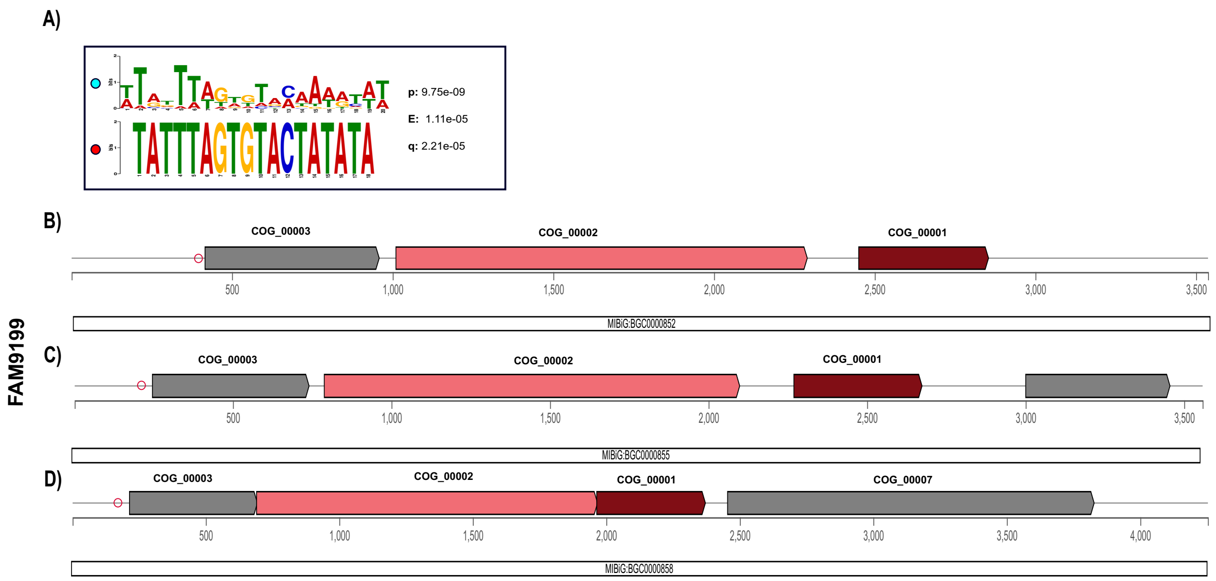


**Figure S5: Recovery of EctR1-mediated regulation in BGC family FAM9199. (A)** TOMTOM comparison of the EctR1 binding site (TATTTAGTGTACTATATA) against the predicted motif upstream of COG_00003 (*ectA*) in FAM9199 (p, p-value; e, E-value; q, q-value) ; coloured circles indicate motif source (blue, pipeline-predicted; red, literature reference). **(B–D)** Genomic architecture of the three MIBiG anchors BGCs in FAM9199, with COG assignments shown above each gene. **(B)** BGC0000852 (*Sporosarcina pasteurii*) with three-gene cluster encoding COG_00003 (*ectA*), COG_00002 (*ectB*) and COG_00001 (*ectC*). **(C)** BGC0000855 (*Methylomicrobium kenyense*). Same three biosynthetic COGs plus one additional gene (hypothetical protein. No COG formed). **(D)** BGC0000858 (*Methylobacter marinus*) with the same three biosynthetic COGs plus COG_00007 immediately downstream of *ectC*. Gene colours indicate predicted function: burgundy = biosynthetic, pink = biosynthetic-additional, grey = other.

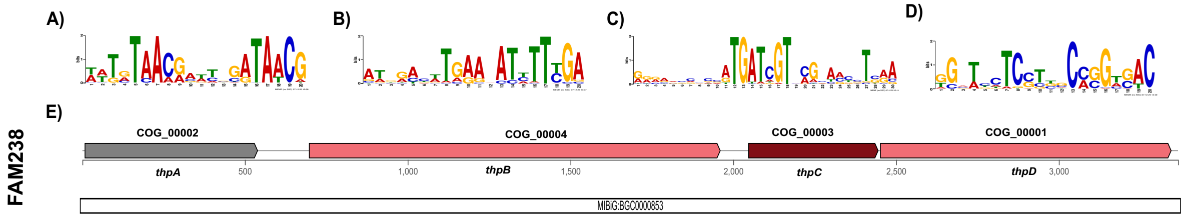


**Supplementary Figure S6: Conserved motifs upstream of the hydroxyectoine biosynthetic genes in FAM238. (A–D)** Sequence logo of the most significant motif (lowest MEME E-value) upstream of each biosynthetic COG: COG_00002/*thpA* (A), COG_00004/*thpB* (B), COG_00003/*thpC* (C) and COG_00001/*thpD* (D); all four have $E < 10^{-116}$. None matched the EctR1 or GlnR reference sites**. (E)** Genomic architecture of the MIBiG anchor BGC0000853 (*Streptomyces anulatus*), genes coloured by predicted function (burgundy, biosynthetic; pink, biosynthetic-additional; grey, other).

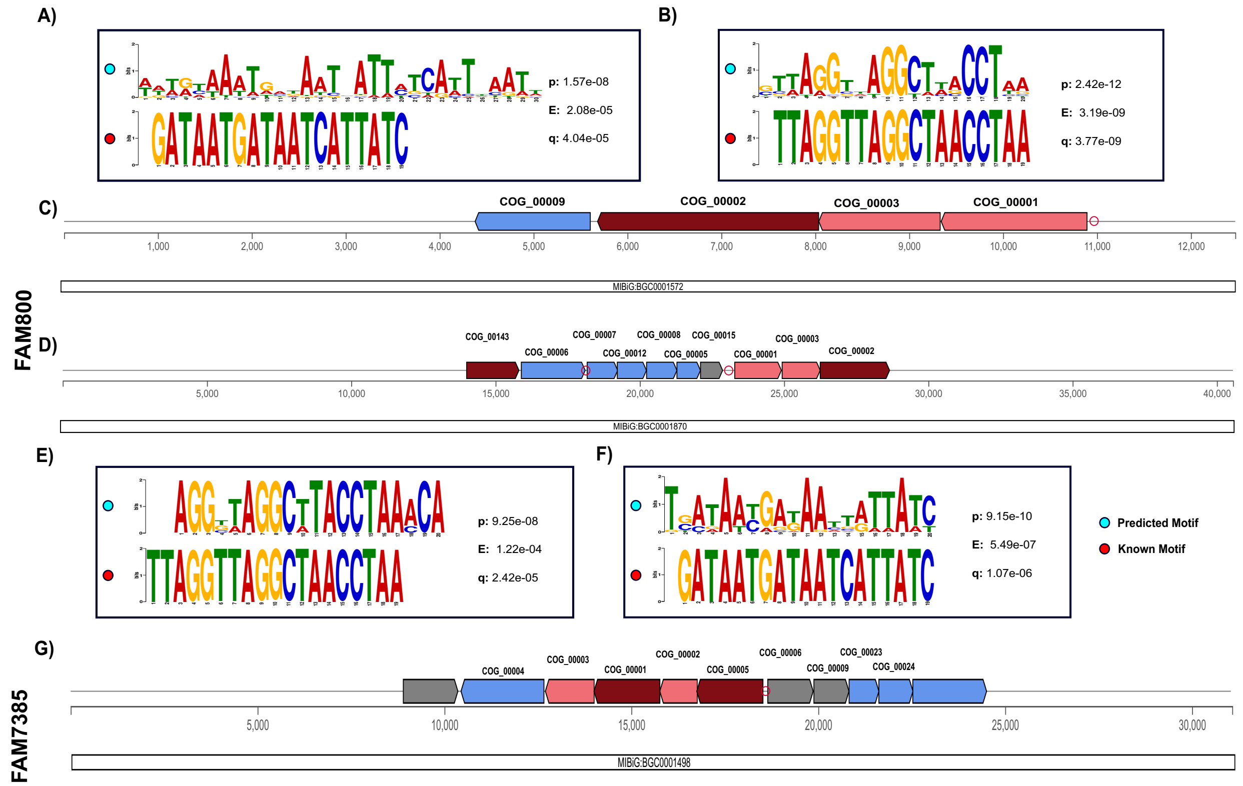


**Figure S7: TOMTOM recovery of predicted regulatory motifs in siderophore BGC families FAM800 and FAM7385.** Pairwise TOMTOM comparisons of pipeline-predicted motifs upstream of COGs in FAM800 (A, B and E) and FAM7385 (F) against known iron-responsive regulator binding sites. In each comparison, the top logo represents the pipeline-predicted motif and the bottom logo the reference binding site; coloured circles indicate motif source (blue, pipeline-predicted; red, literature reference). **(A)** TOMTOM comparison of COG_00001 motif_2 (z30) against the Fur consensus (p, p-value; e, E-value; q, q-value). **(B)** TOMTOM comparison of COG_00001 motif_1 (z20) against the IdeR operator, the iron box recognised by DtxR-family regulators including DmdR1. **(C)** Genomic architecture of the MIBiG anchor BGC0001572 (desferrioxamine E, *Pantoea agglomerans*). **(D)** Genomic architecture of BGC0001870 (putrebactin, *Xenorhabdus budapestensis*). **(E)** TOMTOM comparison of COG_00007 motif_1 against the IdeR consensus. (F) TOMTOM comparison of COG_00005 motif_4 against the Fur box consensus. **(G)** Genomic architecture of the MIBiG anchor BGC0001498 (aerobactin, *Xenorhabdus szentirmaii* DSM 16338). Gene colours indicate predicted function (burgundy, biosynthetic; pink, biosynthetic-additional; blue, transport; green, regulatory; grey, other).

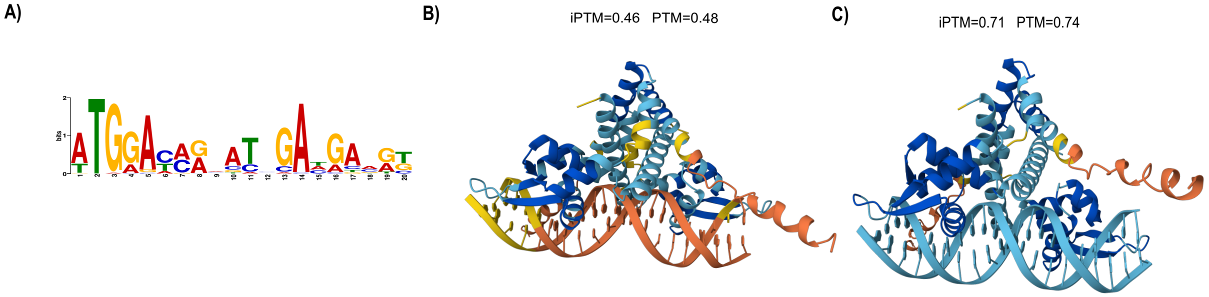


**Figure S8: AlphaFold3 models of the MarR regulator of FAM9025 (BGC1007481, *Teredinibacter sp.*) with its two upstream motifs.** Three significant motifs were recovered for this cluster under the zoops_maxw20 setting: the *ectA* operator (COG_00004 motif_1, 165 bp upstream; Figure 2C) and two motifs upstream of the MarR gene COG_00007, motif_5 at 91 bp (E = $3.7\times10^{-43}$; Figure 2D) and motif_1 at 1 bp from the start codon. TOMTOM matched the *ectA* and motif_5 sites to the EctR1 operator. (A) Sequence logo of the promoter-proximal COG_00007 motif_1 (E = $2.4\times10^{-85}$). Its position is consistent with MarR-family autorepression, in which the regulator binds at or near its own transcription start (REF). (B–C) Dimer–DNA models for COG_00007 motif_5 (B; ipTM 0.46 vs 0.50 for a composition-matched scrambled control) and COG_00007 motif_1 (C; 0.71 vs 0.49), coloured by pLDDT; grey arrows link each motif to its position relative to the gene. The ectA complex is shown in Figure 2E; all scores are in Table S2.

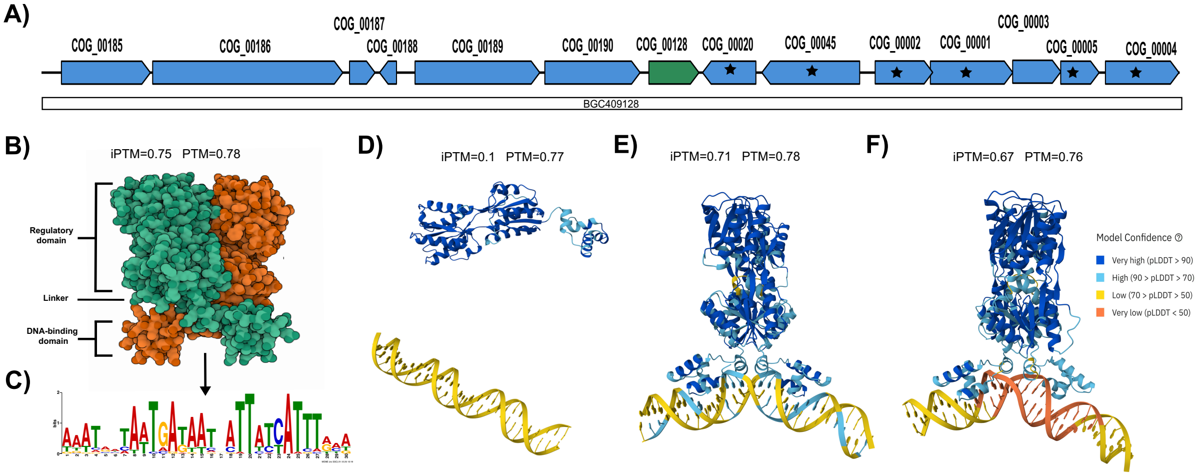


**Figure S9: AlphaFold3 models of the GalR/LacI regulator of FAM123604 (BGC409128, *Aeromonas sp. HMWF016*). (A)** Genomic architecture of BGC409128; the GalR/LacI TF (COG_00128) is green, and starred genes carry COG_00002 motif_1 upstream. **(B)** Homodimer model (ipTM = 0.75, pTM = 0.78), consistent with the LacI-family quaternary structure (REF). **(C)** Sequence logo of COG_00002 motif_1 (E = 2×10$^{-101}$), the most significant motif in the family, which also occurs upstream of the TF gene itself (FIMO match). **(D)** Monomer–DNA model for motif_1; protein and DNA remain dissociated (ipTM = 0.10, real and scrambled). **(E)** Dimer–DNA model for motif_1 (ipTM = 0.70), in the expected LacI-type geometry. **(F)** Dimer–DNA model for the composition-matched scrambled motif_1 control (ipTM = 0.67). The near-identical scores in (E) and (F) indicate a structurally plausible but not sequence-specific interaction. Protein and DNA coloured by pLDDT; scores in **Table S3**.

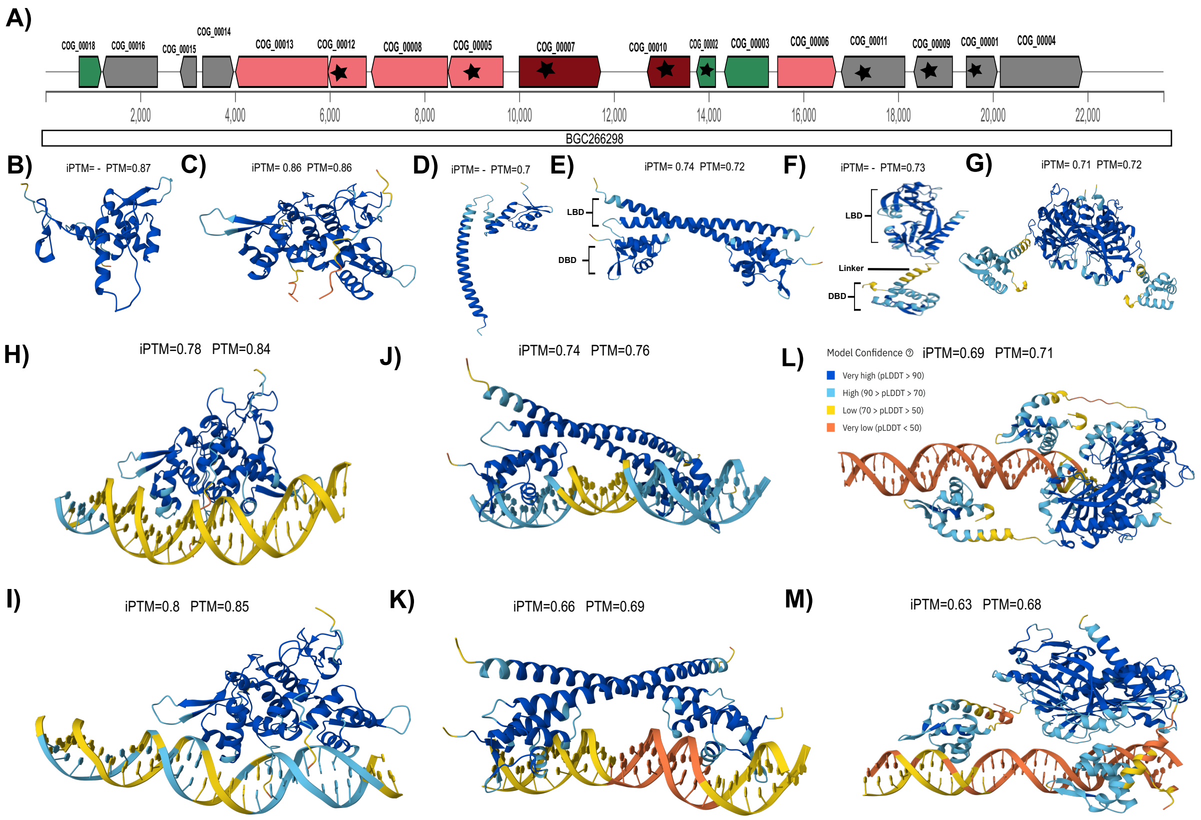


**Figure S10: AlphaFold3 models of the three regulators of FAM2960 (BGC266298, *Pseudomonas_E prosekii*). (A)** Genomic architecture of BGC266298, genes coloured by predicted function (burgundy, biosynthetic; pink, biosynthetic-additional; green, regulatory; grey, other), COG assignments above each gene; starred genes carry the shared COG_00010 motifs upstream. **(B–G)** Monomer and dimer models of MerR (COG_00002; B, C), LysR (COG_00003; D, E) and HxlR (COG_00018; F, G); all three fold and dimerise confidently (**Table S4**). **(H–M)** Dimer–DNA models against COG_00010 motif_2 and motif_3, which occur upstream of eight COGs in FAM2960, for HxlR (H, I), MerR (J, K) and LysR (L, M); scores in **Table S5**. HxlR scores highest but also docks confidently onto every one of nine unrelated motifs (**Table S6**), so its scores reflect fold rather than sequence recognition. No scrambled controls were run for this family. Protein and DNA coloured by pLDDT.

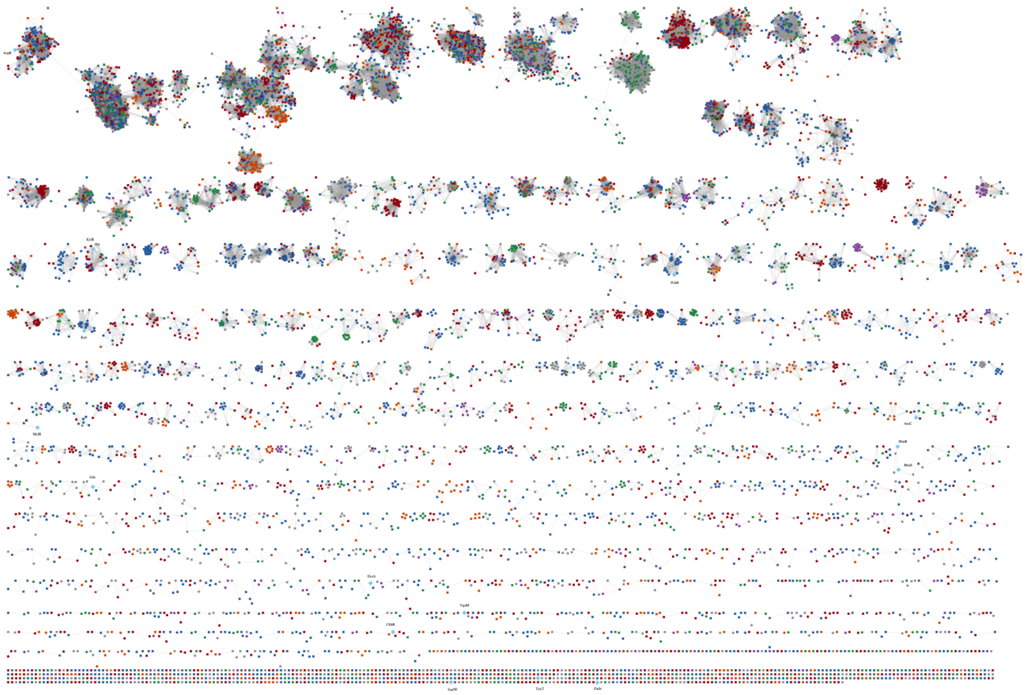

**Figure S11:** Sequence similarity network of all AraC/XylS-annotated TF sequences. Each node represents a unique TF sequence, and each edge represents pairwise sequence similarity at an alignment score threshold of 45. Nodes are coloured by the BGC class of the source BGC: orange = RiPP, blue = NRP, green = Polyketide, purple = Terpene, red = Other, grey = hybrid class. Larger light blue diamonds indicate PRODORIC-validated TFs incorporated as validation anchors.

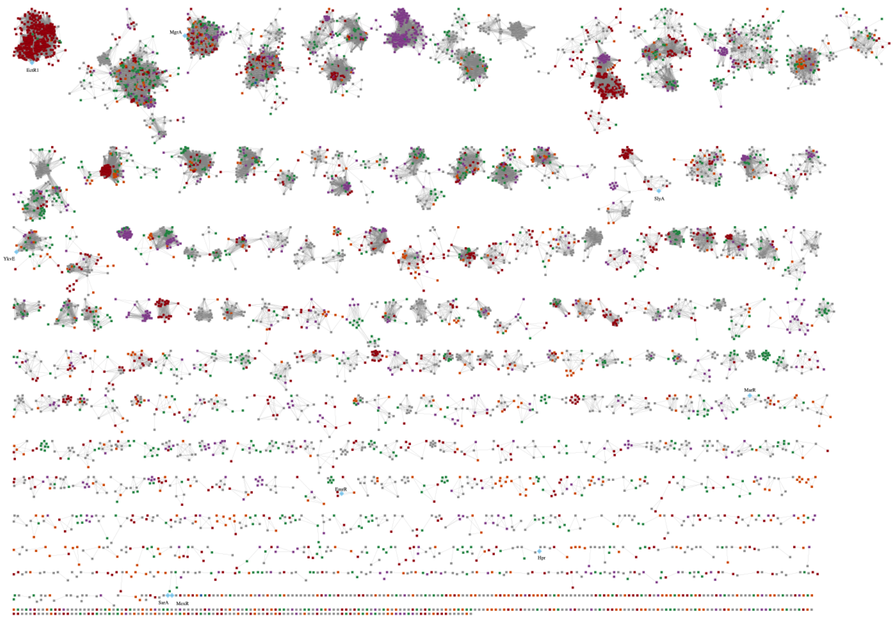

**Figure S12: Sequence similarity network of all MarR-annotated TF sequences.** Each node represents a unique TF sequence, and each edge represents pairwise sequence similarity at an alignment score threshold of 24. Nodes are coloured by the BGC class of the source BGC: orange = RiPP, blue = NRP, green = Polyketide, purple = Terpene, red = Other, grey = hybrid class. Larger light blue diamonds indicate PRODORIC-validated TFs incorporated as validation anchors.

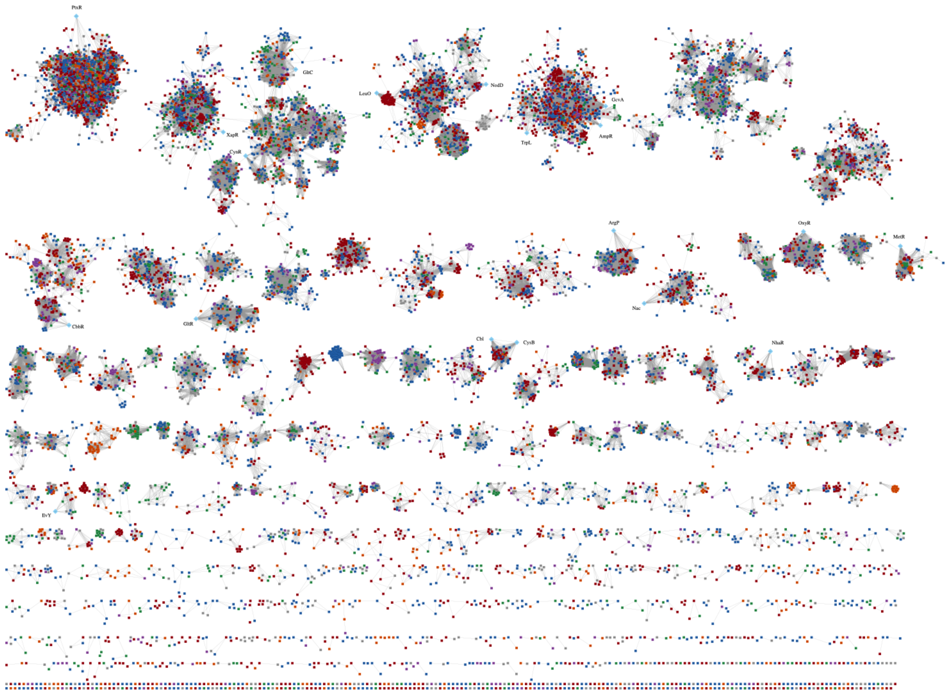

**Figure S13: Sequence similarity network of all LysR-annotated TF sequences.** Each node represents a unique TF sequence, and each edge represents pairwise sequence similarity at an alignment score threshold of 45. Nodes are coloured by the BGC class of the source BGC: orange = RiPP, blue = NRP, green = Polyketide, purple = Terpene, red = Other, grey = hybrid class. Larger light blue diamonds indicate PRODORIC-validated TFs incorporated as validation anchors.

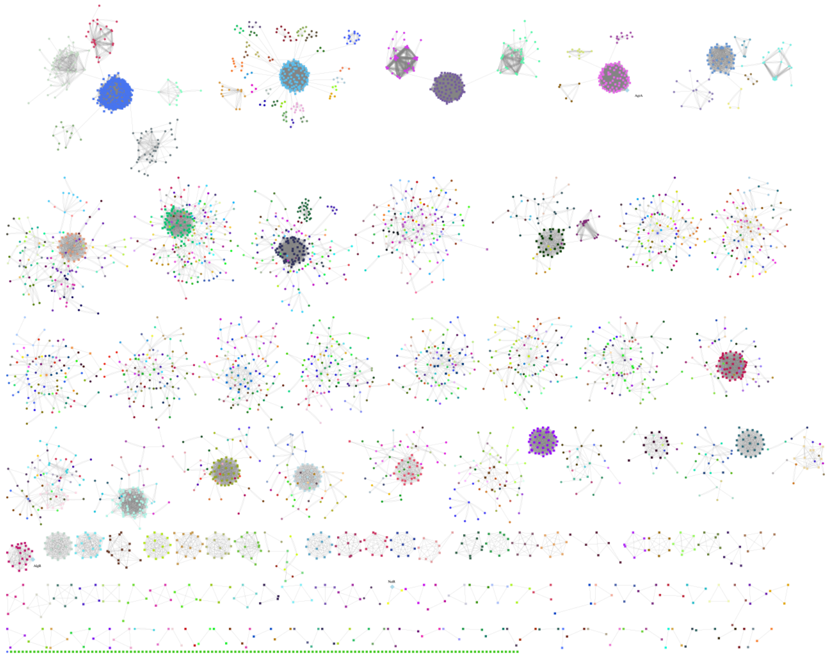

**Figure S14: Combined TF and motif network for the LytTR family.** Each rectangle represents a unique TF sequence, coloured by SSN cluster. Each ellipse represents an individual upstream motif, coloured by motif MCL cluster. TF nodes are connected to one representative motif node per motif cluster, and TF nodes within the same SSN cluster are connected by pairwise sequence similarity edges (weight = alignment score). Motif nodes within the same motif cluster are connected by TOMTOM similarity edges (weight = −log10 q-value). Only motif clusters with family coverage ≥ 0.5 are shown. Larger light blue diamonds indicate PRODORIC-validated TFs.

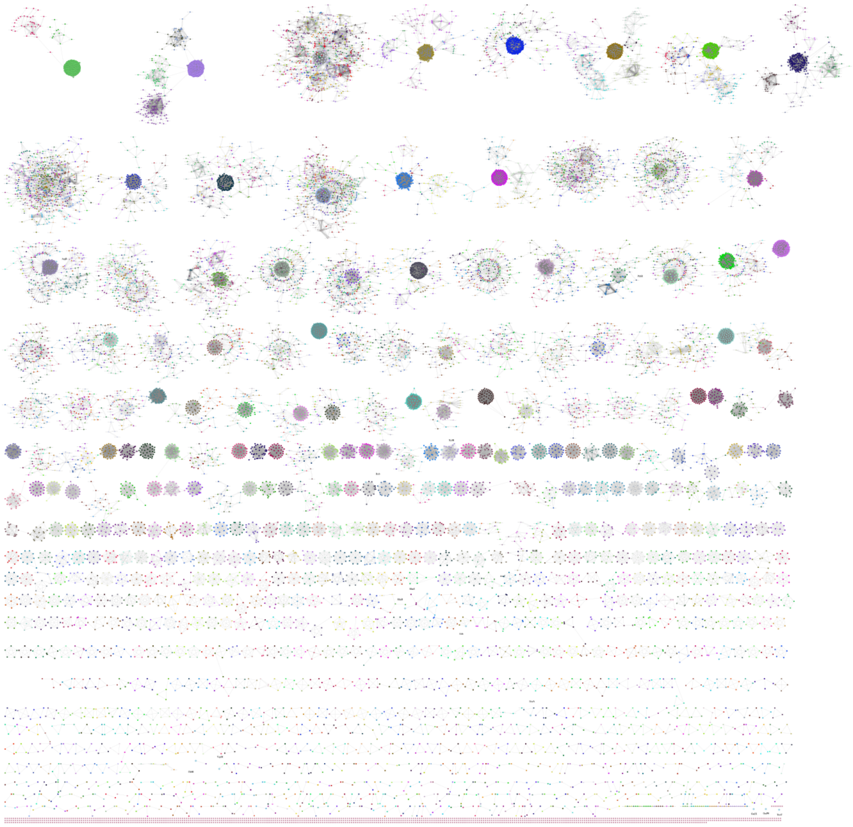

**Figure S15: Combined TF and motif network for the AraC/XylS family.** Each rectangle represents a unique TF sequence, coloured by SSN cluster. Each ellipse represents an individual upstream motif, coloured by motif MCL cluster. TF nodes are connected to one representative motif node per motif cluster, and TF nodes within the same SSN cluster are connected by pairwise sequence similarity edges (weight = alignment score). Motif nodes within the same motif cluster are connected by TOMTOM similarity edges (weight = −log10 q-value). Only motif clusters with family coverage ≥ 0.5 are shown. Larger light blue diamonds indicate PRODORIC-validated TFs.

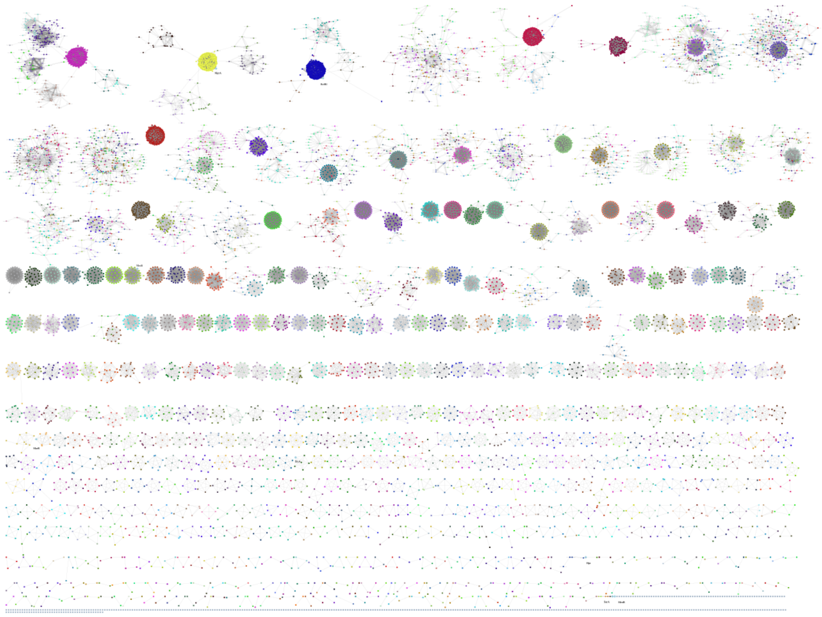

**Figure S16: Combined TF and motif network for the MarR family.** Each rectangle represents a unique TF sequence, coloured by SSN cluster. Each ellipse represents an individual upstream motif, coloured by motif MCL cluster. TF nodes are connected to one representative motif node per motif cluster, and TF nodes within the same SSN cluster are connected by pairwise sequence similarity edges (weight = alignment score). Motif nodes within the same motif cluster are connected by TOMTOM similarity edges (weight = −log10 q-value). Only motif clusters with family coverage ≥ 0.5 are shown. Larger light blue diamonds indicate PRODORIC-validated TFs.

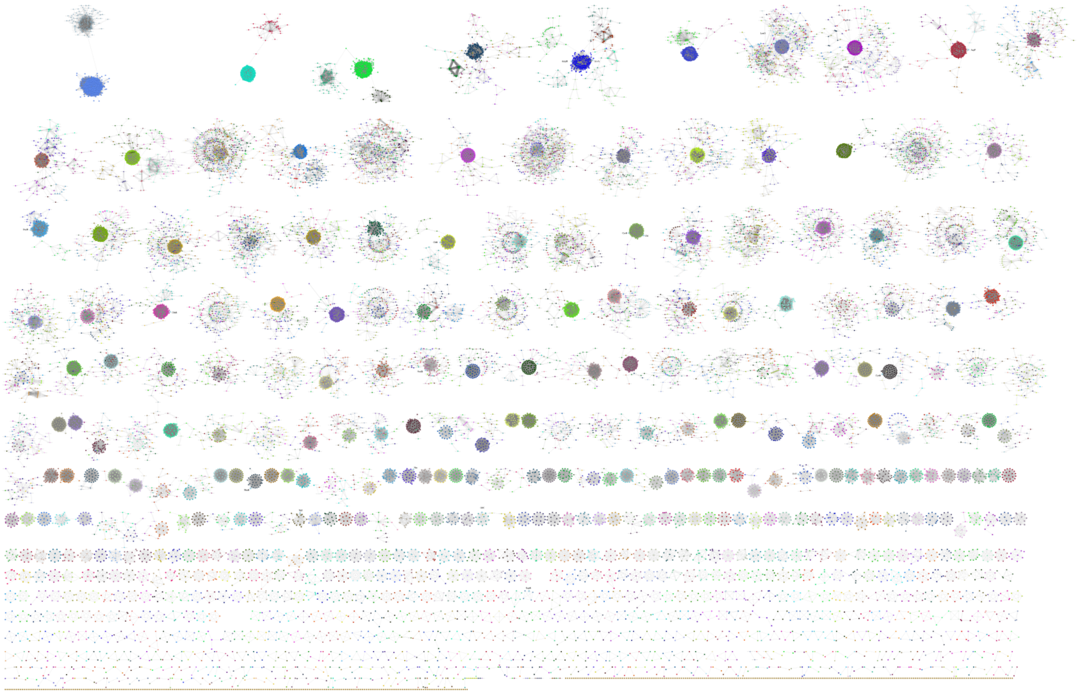

**Figure S17: Combined TF and motif network for the LysR family.** Each rectangle represents a unique TF sequence, coloured by SSN cluster. Each ellipse represents an individual upstream motif, coloured by motif MCL cluster. TF nodes are connected to one representative motif node per motif cluster, and TF nodes within the same SSN cluster are connected by pairwise sequence similarity edges (weight = alignment score). Motif nodes within the same motif cluster are connected by TOMTOM similarity edges (weight = −log10 q-value). Only motif clusters with family coverage ≥ 0.5 are shown. Larger light blue diamond’s indicate PRODORIC-validated TFs.

#### Supplementary Table:

**Table S1: Sequence similarity network parameters for the 20 PSTF families analysed.** For each TF family, pipeline-annotated TF sequences (deduplicated at 90% identity) were combined with PRODORIC-characterised regulators of the same family and submitted to EFI-EST. Sequences outside the length range shown were excluded before network construction. The alignment-score threshold was chosen from the percent-identity versus alignment-score plot of each network. SSN TF clusters were defined by MCL clustering (inflation 2.0); singletons are not counted.

| TF family | Min. length (aa) | Max. length (aa) | Alignment-score threshold | SSN TF clusters |
|---|---|---|---|---|
| LysR | 270 | 350 | 45 | 890 |
| AraC/XylS | 190 | 415 | 45 | 1,006 |
| LytTR | 215 | 310 | 40 | 133 |
| MarR | 120 | 220 | 24 | 645 |
| ArsR | 85 | 145 | 15 | 65 |
| AsnC | 130 | 180 | 30 | 81 |
| Crp | 170 | 480 | 40 | 261 |
| DeoR | 240 | 270 | 40 | 66 |

| | | | | |
|---|---|---|---|---|
| Fis | 425 | 520 | 50 | 14 |
| GalR/LacI | 300 | 400 | 50 | 286 |
| GerE | 180 | 280 | 45 | 734 |
| GntR | 190 | 290 | 55 | 839 |
| Histone-like DNA-binding | 80 | 120 | 17 | 14 |
| HxlR | 100 | 180 | 20 | 44 |
| MerR | 120 | 300 | 50 | 521 |
| PadR | 105 | 220 | 20 | 78 |
| ROK | 290 | 420 | 40 | 115 |
| SinR | 170 | 225 | 25 | 65 |
| TetR/AcrR | 170 | 270 | 30 | 1,925 |
| Trans_reg_C | 110 | 280 | 80 | 873 |

156

**Table S2: AlphaFold3 scores for the MarR regulator of BGC1007481 (FAM9025) modelled against three predicted motifs.** Each motif was supplied as double-stranded DNA and modelled in complex with the TF as a monomer and as a homodimer, alongside a composition-matched control generated by shuffling the motif sequence (fixed seed). pTM estimates overall fold accuracy (> 0.5, fold likely correct); ipTM estimates the accuracy of subunit positioning within a complex (> 0.8 confident, 0.6–0.8 uncertain, < 0.6 likely failed). Protein-only models: monomer pTM 0.80; dimer ipTM 0.76, pTM 0.78. For the ectA motif the monomer distinguishes real from scrambled DNA (0.47 vs 0.10) while the dimer does not (0.75 for both), because the protein–protein interface dominates dimer ipTM; for the promoter-proximal motif_1 only the dimer shows a difference (0.71 vs 0.49); for motif_5 neither does. Each comparison rests on a single model pair.

| Run | Monomer real ipTM | Monomer scrambled ipTM | Dimer real ipTM | Dimer scrambled ipTM |
|---|---|---|---|---|
| **COG_00004 motif_1 (*ectA*)** | 0.47 | 0.10 | 0.75 | 0.75 |
| **COG_00007 motif_1 (promoter-proximal)** | 0.11 | 0.10 | 0.71 | 0.49 |
| **COG_00007 motif_5**<br><br>**(candidate autorgulatory)** | 0.09 | 0.10 | 0.46 | 0.50 |

**Table S3: AlphaFold3 confidence scores for the GalR/LacI regulator of BGC409128 (*Aeromonas* sp. HMWF016) modelled against the family regulon motif.** The TF was modelled as a monomer and a homodimer, both alone (protein only) and in complex with the predicted regulon motif (motif_1, upstream of COG_00002) and a composition-matched scrambled control. ipTM (interface predicted TM-score) reports confidence in the relative positioning of subunits within a complex and is the primary readout of TF–DNA interaction; pTM reports confidence in the overall fold. ipTM is undefined for the protein-only monomer (single chain, no interface). Values ≥0.8 indicate confident predictions, ≤0.6 likely failed predictions, and 0.6–0.8 a grey zone. The monomer did not engage DNA (ipTM = 0.10, real and scrambled), while the dimer formed a complex with the motif (ipTM = 0.70) but scored near-identically against the scrambled control (ipTM = 0.67), indicating a structurally plausible but not sequence-specific interaction.

| **Run** | **ipTM** | **pTM** |
|---|---|---|
| Monomer (protein only) | — | 0.88 |
| Dimer (protein only) | 0.75 | 0.78 |
| Monomer vs motif_1 | 0.10 | 0.77 |
| Monomer vs motif_1 (scrambled) | 0.10 | 0.77 |
| Dimer vs motif_1 | 0.70 | 0.78 |
| Dimer vs motif_1 (scrambled) | 0.67 | 0.76 |

**Table S4: AlphaFold3 fold and oligomerisation confidence for the three co-encoded TFs of BGC266298 (*Pseudomonas_E prosekii*).** Each TF was modelled as a monomer and a homodimer to confirm its oligomeric state prior to TF–DNA complex modelling. Monomer pTM reports overall fold confidence; dimer ipTM and pTM report confidence in the subunit interface and overall dimer fold, respectively. All three TFs formed confident folds and homodimers, consistent with the known functional states of the MerR, LysR and HxlR families, justifying the use of the dimer as the structural unit in subsequent DNA-complex modelling. Panel references correspond to Figure S9.

| **TF (COG)** | **Family** | **Monomer pTM** | **Dimer ipTM** | **Dimer pTM** | **Figure** |
|---|---|---|---|---|---|
| COG_00002 | MerR | 0.70 | 0.74 | 0.72 | S9B, S9C |

| | | | | | |
|---|---|---|---|---|---|
| COG_00003 | LysR | 0.73 | 0.71 | 0.72 | S9D, S9E |
| COG_00018 | HxlR | 0.87 | 0.86 | 0.86 | S9F, S9G |

**Table S5: Dimer–DNA AlphaFold3 ipTM scores for the three BGC266298 TFs against the two shared regulon motifs.** Each TF homodimer was modelled in complex with the two top-ranked regulon motifs (motif_2 and motif_3, both upstream of COG_00010), which are conserved across eight COGs within FAM2960. ipTM ≥0.8 indicates confident binding, ≤0.6 a likely failed prediction, and 0.6–0.8 a grey zone. All corresponding monomer–DNA predictions scored ipTM <0.4 and are not shown.

| **TF (COG)** | **motif_2** | **motif_3** |
|---|---|---|
| HxlR (COG_00018) | 0.78 | 0.80 |
| MerR (COG_00002) | 0.74 | 0.66 |
| LysR (COG_00003) | 0.69 | 0.63 |

**Table S6: Dimer–DNA AlphaFold3 ipTM scores for the three BGC266298 TFs against the two shared regulon motifs.** Each TF homodimer was modelled in complex with the two top-ranked regulon motifs (motif_2 and motif_3, both upstream of COG_00010), which are conserved across eight COGs within FAM2960. ipTM ≥0.8 indicates confident binding, ≤0.6 a likely failed prediction, and 0.6–0.8 a grey zone. All corresponding monomer–DNA predictions scored ipTM <0.4 and are not shown. Interpretations are qualitative summaries of the relative scores; scrambled-sequence controls were not run for this dataset, so sequence specificity cannot be confirmed from these values alone.

| **TF dimer** | **COG2 m1** | **COG2 m2** | **COG2 m3** | **COG3 m1** | **COG3 m2** | **COG3 m3** | **COG18 m1** | **COG18 m2** | **COG18 m3** |
|---|---|---|---|---|---|---|---|---|---|
| **MerR** (COG_00002) | 0.69 | 0.68 | 0.67 | 0.64 | 0.68 | 0.68 | 0.66 | 0.75 | 0.66 |

| | | | | | | | | | |
|---|---|---|---|---|---|---|---|---|---|
| **LysR** (COG_00003) | 0.66 | 0.60 | 0.64 | 0.62 | 0.66 | 0.61 | 0.66 | 0.62 | 0.73 |
| **HxlR** (COG_00018) | **0.83** | **0.80** | **0.81** | **0.88** | **0.88** | 0.76 | **0.90** | 0.78 | **0.90** |
